# Mcm10 and RecQL4 Drive the Activation of the Eukaryotic Replicative DNA Helicase CMG

**DOI:** 10.1101/2025.08.05.668837

**Authors:** Riki Terui, Larissa Sambel, Jinho Park, Elyse Hsin-Lien Hwang, Benjamin Duewell, Dan Song, Gheorghe Chistol

**Affiliations:** Chemical and Systems Biology, Stanford School of Medicine, Stanford CA94305; Kyushu University, Stanford School of Medicine, Stanford CA94305, Fukuoka, Japan; Eikon Therapeutics Inc.; Cancer Biology Program, Stanford School of Medicine, Stanford CA94305; Stanford Cancer Institute, Stanford School of Medicine, Stanford CA94305; BioX Interdisciplinary Institute, Stanford School of Medicine, Stanford CA94305

## Abstract

Chromosomes are copied from thousands of origins. At each origin, two replicative DNA helicases are first assembled, then activated to begin unwinding DNA. Several replication proteins are subsequently recruited to the active helicase, forming a replisome. The helicase must undergo dramatic conformational changes during its activation and this process is poorly understood, especially in metazoa. How the metazoan replicative helicase is activated, and which proteins promote this essential process are long-standing questions. Using a combination of single-molecule imaging and ensemble biochemistry, we show that Mcm10 and RecQL4 act in a concerted manner to activate replicative helicases. Mcm10 first binds to inactive helicases, then recruits RecQL4, which synergizes with Mcm10 to promote helicase activation. Mcm10 is not incorporated into replisomes and dissociates from origins during replication initiation. In the absence of Mcm10, RecQL4 is recruited to origins via an interaction with the Mcm7 subunit of the helicase. Our data reveal that Mcm10 and RecQL4 play partially redundant roles during helicase activation, help resolve long-standing controversies about the roles of Mcm10 and RecQL4 in DNA replication, and reveal replication initiation defects caused by pathologic RecQL4 mutations.

## INTRODUCTION

During cell division, eukaryotic genomes are replicated in parallel from thousands of replication origins. Each replication origin fires bidirectionally, ensuring that chromosomes are fully replicated regardless of the exact number or location of individual origins. Replication initiation is strictly regulated to maintain genome integrity, and dysregulation of origin firing is linked to genomic instability and replication stress, as reported in many cancers^1–4^. Despite its importance in maintaining genome integrity, replication initiation remains poorly understood, especially in metazoa.

Origins are licensed for replication during late mitosis and G1, when two Mcm2-7 complexes (Mini Chromosome Maintenance proteins 2-7) are loaded onto double-stranded DNA (dsDNA), forming a pre-Replication Complex (preRC)^5,6^. During S-phase, two copies of Cdc45 (Cell Division Control protein 45) and two copies of GINS (Go-Ichi-Ni-San, Japanese for 5-1-2-3, referring to subunits Sld5, Psf1, Psf2, and Psf3) are recruited to the preRC^7^. This completes the assembly of two replicative helicases, also known as CMG (Cdc45•Mcm2-7•GINS)^8^. Cdc45 and GINS recruitment is promoted by DDK (Dbf4-Dependent Kinase) and CDK2 (Cyclin-Dependent Kinase 2)^7^. In metazoa, several firing factors facilitate CMG assembly, including TopBP1 (Topoisomerase 2 Binding Protein 1), TRESLIN (TOPBP1-interacting, Replication-Stimulating protein), MTBP (MDM2 Binding Protein), and DONSON (protein Downstream Neighbor of SON)^9–15^. After assembly, the two CMGs remain in an inactive head-to-head configuration and encircle dsDNA. Two copies of DONSON are tightly bound to this inactive CMG dimer^16,17^. We refer to this (CMG•DONSON)_2_ complex as the “pre-activation complex” (Fig. 1a).

**Figure 1:**
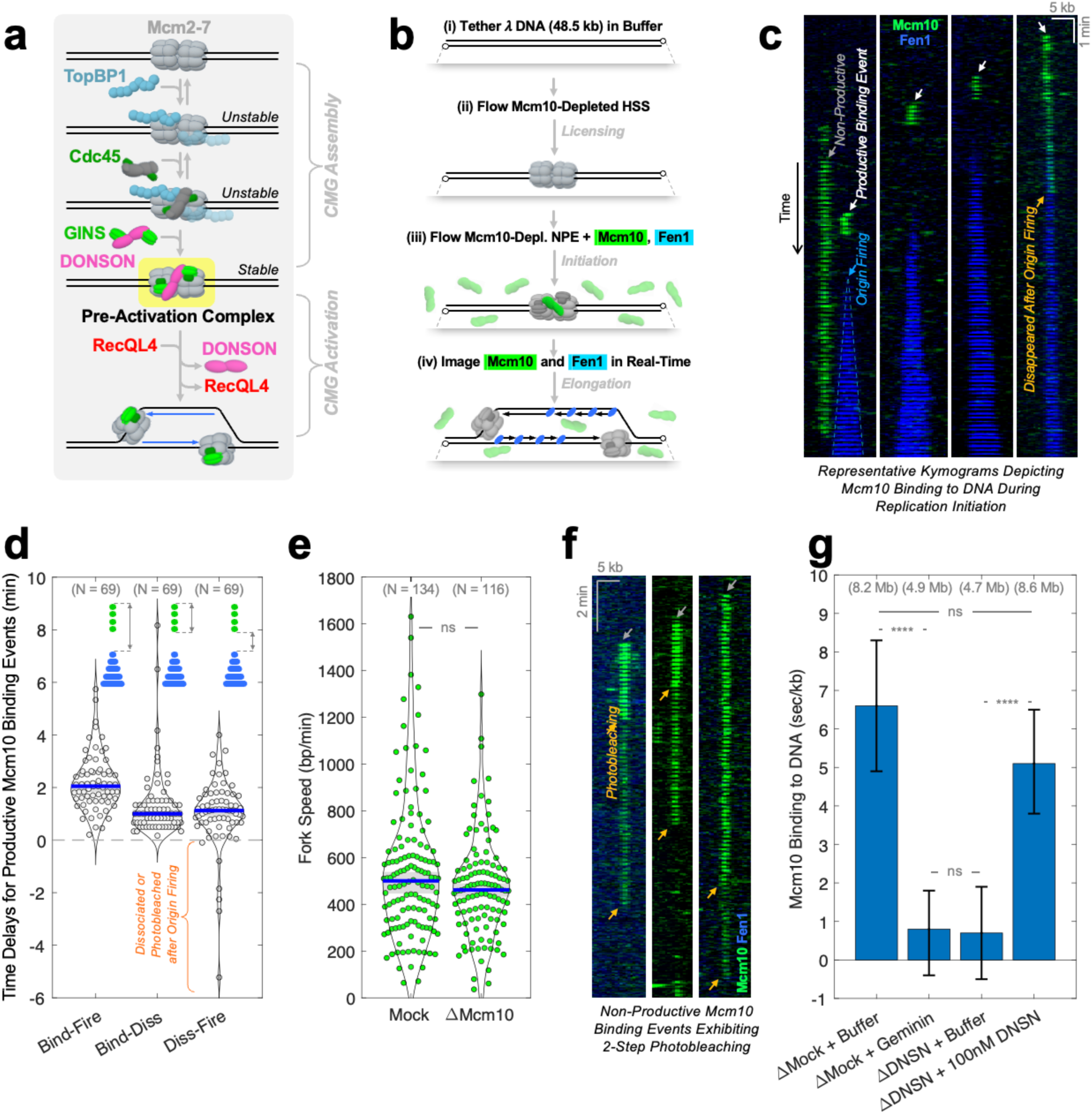
**(a)** Current model of replication initiation in metazoa. **(b)** Workflow of the single-molecule experiment to visualize Mcm10^AF647^ (green) binding to DNA, Fen1^mKikGR^ (blue) acts as a marker of nascent DNA. **(c)** Representative kymograms illustrating Mcm10^AF647^ (green) binding to licensed DNA. Fen1^mKikGR^ (blue) marks nascent DNA. **(d)** Time delays for productive Mcm10^AF647^ binding events. Left: delay between Mcm10^AF647^ binding and origin firing. Center: delay between Mcm10^AF647^ binding and dissociation. Right: delay between Mcm10^AF647^ dissociation and origin firing (negative values mean that Mcm10^AF647^ dissociated/photobleached after origin firing). N represents the number of productive Mcm10^AF647^ binding events. **(e)** Fork speed measured in mock-depleted or Mcm10-depleted reactions. N denotes the number of replication forks. **(f)** Representative kymograms of non-productive Mcm10^AF647^ binding events exhibiting two-step photobleaching (orange arrows). **(g)** Quantification of Mcm10^AF647^ binding to DNA normalized to DNA length (Extended Data Fig. 1f illustrates how normalized binding is measured). Geminin blocks licensing and serves as a negative control. Total DNA length analyzed for each condition is reported in parentheses. Error-bars denote the 95% Confidence Interval (CI) estimated via bootstrapping. In **(d)** and **(e)** blue bars represent median values, gray boxes mark the 95% CI. The two-sample Kolmogorov-Smirnov test was used to compute p-values: ns (not significant, p > 0.05), * (p < 0.05), ** (p < 0.01), *** (p < 0.001), **** (p < 0.0001).

To initiate replication, the two CMGs are activated in a multi-step process: DONSON dissociates from CMG^17^, the ATPase/helicase activity of each CMG is upregulated^7^, the two CMGs separate from each other^18,19^, and each CMG transitions from encircling dsDNA to encircling single-stranded DNA (ssDNA). After their activation, the two CMGs start unwinding DNA bidirectionally away from the origin, and several replication proteins are recruited to each CMG to form a multi-subunit complex known as the replisome. To date, the molecular mechanism of CMG helicase activation remains one of the most poorly understood aspects of replication.

The best molecular characterization of CMG activation comes from yeast, where Mcm10 (Mini Chromosome Maintenance protein 10) promotes helicase activation by directly binding CMG, allosterically stimulating its ATPase/helicase activity, and facilitating the cracking of the Mcm2-7 ring ^18–22^. Degrading Mcm10 *in vivo* or omitting Mcm10 from *in vitro* replication reactions strongly suppresses replication initiation and leads to the accumulation of CMGs on chromatin^23–25^.

By contrast, the mechanism of CMG activation in metazoa is poorly understood and remains controversial. In addition to Mcm10, RecQL4 (RecQ Like helicase 4) has been implicated in metazoan CMG activation. Both proteins are poorly conserved between yeast and vertebrates, especially RecQL4 – whose disordered N-terminus is distantly related to the yeast Sld2, while the structured C-terminus contains an ATPase/helicase domain unique to higher eukaryotes^26^ (Extended Data Fig. 1a). It has been challenging to interrogate the role of Mcm10 and RecQL4 in replication initiation because both proteins are involved in other aspects of genome maintenance, including replication elongation, DNA damage response, and DNA repair^27–30^.

There are several seemingly contradictory reports about the involvement of metazoan Mcm10 in replication initiation. For example, one study reported that depleting Mcm10 from *Xenopus* egg extracts strongly inhibited replication initiation^31^, while a more recent publication concluded that Mcm10 is important for replication elongation, but not for initiation^32^. Studies in human cells^33^ and *Xenopus* egg extracts^31^ independently concluded that Mcm10 is required for CMG helicase assembly, in sharp contrast with reports from yeast that Mcm10 participates in CMG activation^23,25,34^. Finally, there is a well-documented interaction between RecQL4 and Mcm10 which is thought to be important for replication initiation^35^. However, the exact purpose of this interaction remains unclear: it was proposed that Mcm10 may recruit RecQL4 to origins^36^ or vice-versa^37^.

Importantly, Mcm10 and RecQL4 are dysregulated in human diseases: both Mcm10^28,38,39^ and RecQL4^30,40–42^ are frequently over-expressed in rapidly proliferating tumors, and their expression levels often correlate with poor patient prognosis. Mcm10 deficiency leads to genomic instability, slower cell proliferation, and impaired cell survival^43–45^. Mutations in RecQL4 are associated with three related but distinct autosomal recessive disorders: Rothmund-Thompson Syndrome (RTS), Baller-Gerold Syndrome (BGS), and RAPADILINO Syndrome (RS)^46,47^. RTS, BGS, and RS patients exhibit skeletal abnormalities, short stature, poikiloderma, and a high predisposition to osteosarcoma^47^. Notably, most patient mutations reside in the C-terminal portion of RecQL4 that is unique to metazoa. Since RecQL4 mutations linked to RTS, BGS, and RS are not embryonically lethal, any replication initiation defect caused by those pathogenic mutations must be relatively mild. Therefore, a highly sensitive replication initiation assay is needed to investigate the molecular basis of RecQL4 syndromes.

To interrogate the molecular mechanism of CMG activation in metazoa and dissect the roles of Mcm10 and RecQL4 in this process, we employed a combination of single-molecule imaging and ensemble biochemistry in *Xenopus* nuclear egg extracts. These extracts contain all the soluble nuclear proteins and efficiently recapitulate DNA replication and other aspects of genome maintenance^48,49^.

By directly imaging fluorescently-labeled Mcm10 molecules on DNA during replication, we found that Mcm10 binds to origins and dissociates from chromatin shortly before origin firing. We found that Mcm10 is not stably incorporated into replisomes, contradicting previous reports that Mcm10 travels with replication forks. Contrary to previous reports that Mcm10 promotes CMG assembly in metazoa^31,33^, our data indicates that Mcm10 participates in CMG activation. We showed that Mcm10 recruits RecQL4 to replication origins, and that RecQL4 promotes Mcm10 dissociation from origins. In the absence of Mcm10, RecQL4 is recruited to origins via a previously unknown interaction between RecQL4 C-terminus and the winged-helix-domain of Mcm7. Finally, we showed that the presence of Mcm10 masks replication initiation defects caused by mutations in RecQL4’s C-terminus, where most pathogenic mutations are found. Our data reveal that Mcm10 and RecQL4 play partially redundant roles during CMG helicase activation, help resolve long-standing controversies about the function of Mcm10/RecQL4 in metazoan replication initiation and provide valuable mechanistic insights for researchers studying RecQL4 syndromes.

## RESULTS

### Mcm10 Binds to and Dissociates from Replication Origins Before Origin Firing

To directly establish Mcm10’s role in metazoan replication initiation, we visualized its recruitment during DNA replication in *Xenopus* egg extracts. Extract was used to replicate a linear λ DNA substrate stretched to ∼85% of its contour length and immobilized via biotin-streptavidin linkages in a microfluidic flow cell as previously described (Fig. 1bi)^17,50,51^. DNA was licensed for replication via incubation in High-Speed Supernatant (HSS), an extract that mimics G1 (Fig. 1bii). Replication was initiated by adding Nucleo-Plasmic Extract (NPE), an extract that mimics S-phase (Fig. 1biii)^48^. Mcm10-depleted extract was supplemented with 10 nM of recombinant Mcm10 labeled with Alexa Fluor 647 (Mcm10^AF647^). An ensemble plasmid replication assay was used to verify that recombinant Mcm10 was functional (Extended Data Fig.1b-e). A total internal reflection fluorescence microscope was used to visualize the recruitment of individual Mcm10^AF647^ molecules to replicating DNA (Fig. 1biv). We previously used this approach to visualize the dynamics of GINS, Cdc45, RecQL4, TopBP1, and DONSON during replication initiation and elongation^17,51^. To monitor DNA synthesis, we supplemented extract with recombinant Fen1 fused to the fluorescent protein mKikGR (Fen1^mKikGR^) as described previously (Fig. 1biii)^52^. Fen1^mKikGR^ is recruited to DNA-bound PCNA, acting as a marker of nascent DNA (Fig. 1biv).

In ∼90% of origin firing events, we detected Mcm10^AF647^ binding shortly before replication initiation (Fig. 1c, white arrows). Mcm10^AF647^ bound to the origin ∼2 min before origin firing, remained bound at the origin for ∼1 min, and dissociated from DNA ∼1 min before origin firing (Fig. 1d). In 63 out of 69 origin firing events, Mcm10^AF647^ dissociated before the appearance of the Fen1^mKikGR^ signal (Fig. 1d). In only 6 out of 69 initiation events, Mcm10^AF647^ remained on DNA after origin firing but disappeared within ∼5 min (due to dissociation or photobleaching). These findings indicate that in *Xenopus* egg extract, Mcm10 participates almost exclusively in replication initiation, and is not stably incorporated into the replisome. Consistent with this idea, replication fork speed was not altered in Mcm10-depleted (ΔMcm10) extract (Fig. 1e).

In addition to productive Mcm10^AF647^ binding events which led to origin firing (Fig. 1c, white arrows), ∼25% of Mcm10^AF647^ binding events were non-productive (Fig. 1c, gray arrow). We attribute these non-productive binding events to failed origin firing attempts. Non-productive Mcm10^AF647^ binding event often exhibited two-step photobleaching (Fig. 1f, orange arrows), indicating that they correspond to a Mcm10 dimer.

### Mcm10 is Recruited to Origins After CMG Assembly

To determine if Mcm10 participates in CMG assembly or CMG activation (Fig. 1a), we used single-molecule imaging to quantify Mcm10^AF647^ recruitment to DNA in mock-depleted or DONSON-depleted extract (Extended Data Fig. 1f). DONSON-mediated recruitment of GINS to Mcm2-7 is the last known step in CMG assembly^17^. Mcm10^AF647^ recruitment to licensed DNA was almost entirely abolished in DONSON-depleted extract, and this defect was robustly rescued by supplementing DONSON-depleted extract with recombinant DONSON (Fig. 1g). This indicates that Mcm10 is recruited to origins downstream of DONSON binding, i.e. after the assembly of the stable (CMG•DONSON)_2_ pre-activation complex.

### RecQL4 Promotes Mcm10 Dissociation from Origins

We recently showed that RecQL4 promotes CMG activation and the accompanying dissociation of DONSON from origins^17^. Here we show that Mcm10 is also involved in CMG activation (Fig. 1). To dissect the order in which Mcm10 and RecQL4 bind to the pre-activation complex, we visualized Mcm10^AF647^ recruitment to DNA in mock-depleted or RecQL4-depleted extract (Fig. 2a).

**Figure 2:**
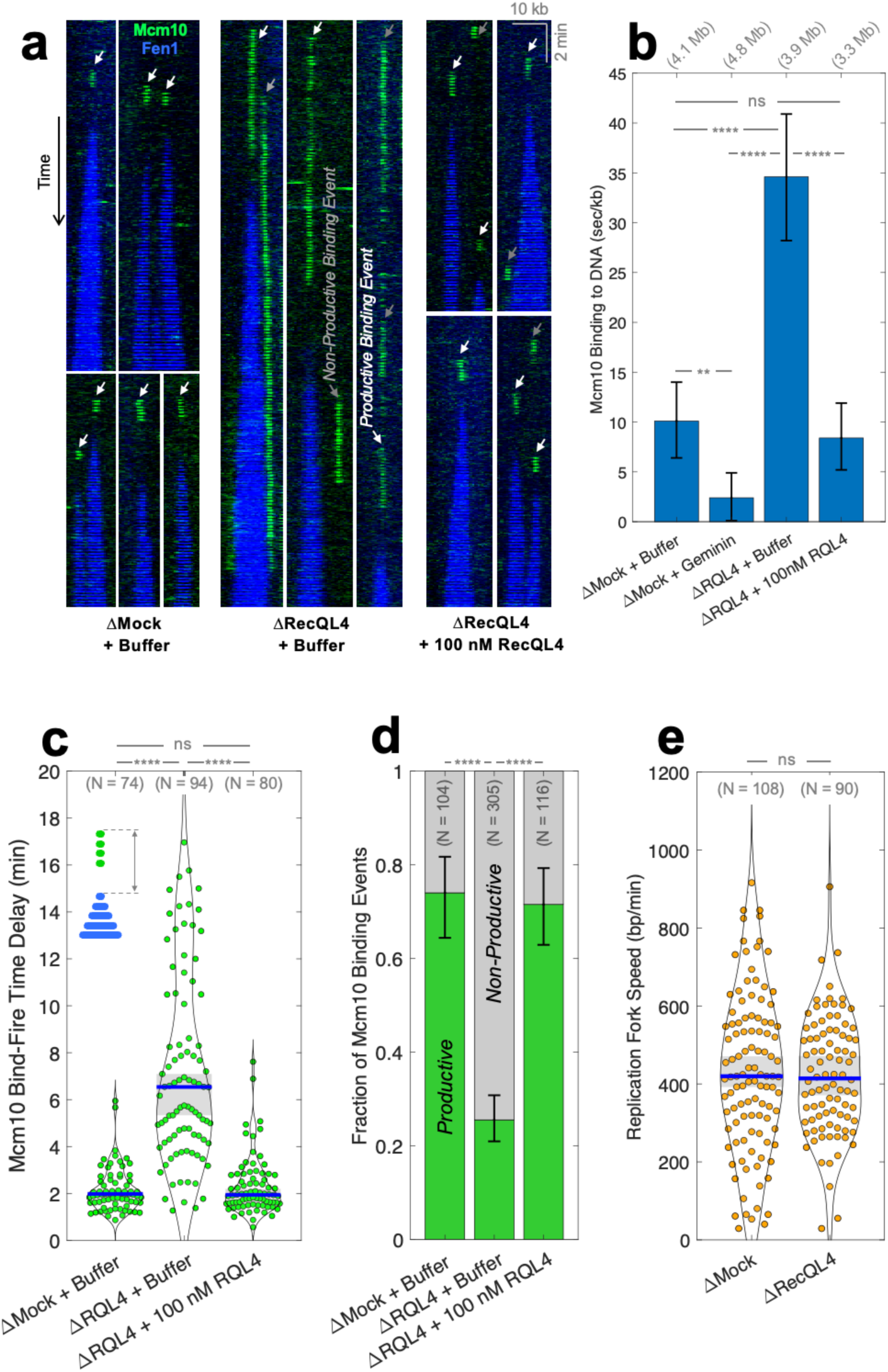
**(a)** Representative kymograms of Mcm10^AF647^ (green) binding licensed DNA. Fen1^mKikGR^ (blue) marks nascent DNA. **(b)** Quantification of Mcm10^AF647^ binding to DNA. Total DNA length analyzed is reported in parentheses. **(c)** Time delay between Mcm10^AF647^ binding and origin firing. N – number of productive events analyzed. Blue bars and gray boxes denote the median and 95% CI. See Extended Data Fig. 2a-b for binding-dissociation and dissociation-firing delays. **(d)** Fraction of productive (green) versus non-productive (gray) Mcm10^AF647^ binding events. N – number of binding events analyzed. See Extended Data Fig. 2c for non-productive event durations. In **(b)** and **(d)** error-bars denote the 95% CI. Two-sample Kolmogorov-Smirnov test was used to compute p-values: ns (not significant, p > 0.05), * (p < 0.05), ** (p < 0.01), *** (p < 0.001), **** (p < 0.0001).

Depleting RecQL4 caused a large reduction in origin firing efficiency, in agreement with our recent report^17^. Mcm10^AF647^ binding to chromatin increased ∼4-fold in RecQL4-depleted extract (Fig. 2b). At origins that did fire in RecQL4-depleted extract, Mcm10^AF647^ resided for ∼6 minutes, 3-fold longer than in mock-depleted extract (Fig. 2c, also see Extended Data Fig. 2a-c). Moreover, in RecQL4-depleted extract, most Mcm10^AF647^ binding events (∼75%) did not result in origin firing (Fig. 2d). Importantly, these replication initiation defects were robustly rescued by adding recombinant RecQL4 to depleted extract (Fig. 2b-d). These findings indicate that RecQL4 normally promotes Mcm10 dissociation from origins.

### Mcm10 Promotes Efficient RecQL4 Binding to Origins

It was previously reported that Mcm10 can interact with RecQL4^35^, and we independently verified that recombinant RecQL4 and Mcm10 interact *in vitro* (Extended Data Fig. 3a-b). However, the exact role of this interaction remains unclear. To address this question, RecQL4-depleted extract was supplemented with 10 nM of recombinant RecQL4^AF647^, and single-molecule imaging was used to monitor its binding to DNA in mock-depleted versus Mcm10-depleted extract (Fig. 3a). RecQL4^AF647^ binding to DNA in Mcm10-depleted extract was ∼4-fold lower than in mock (Fig. 3b), suggesting that Mcm10 helps recruit RecQL4 to the pre-activation complex.

**Figure 3:**
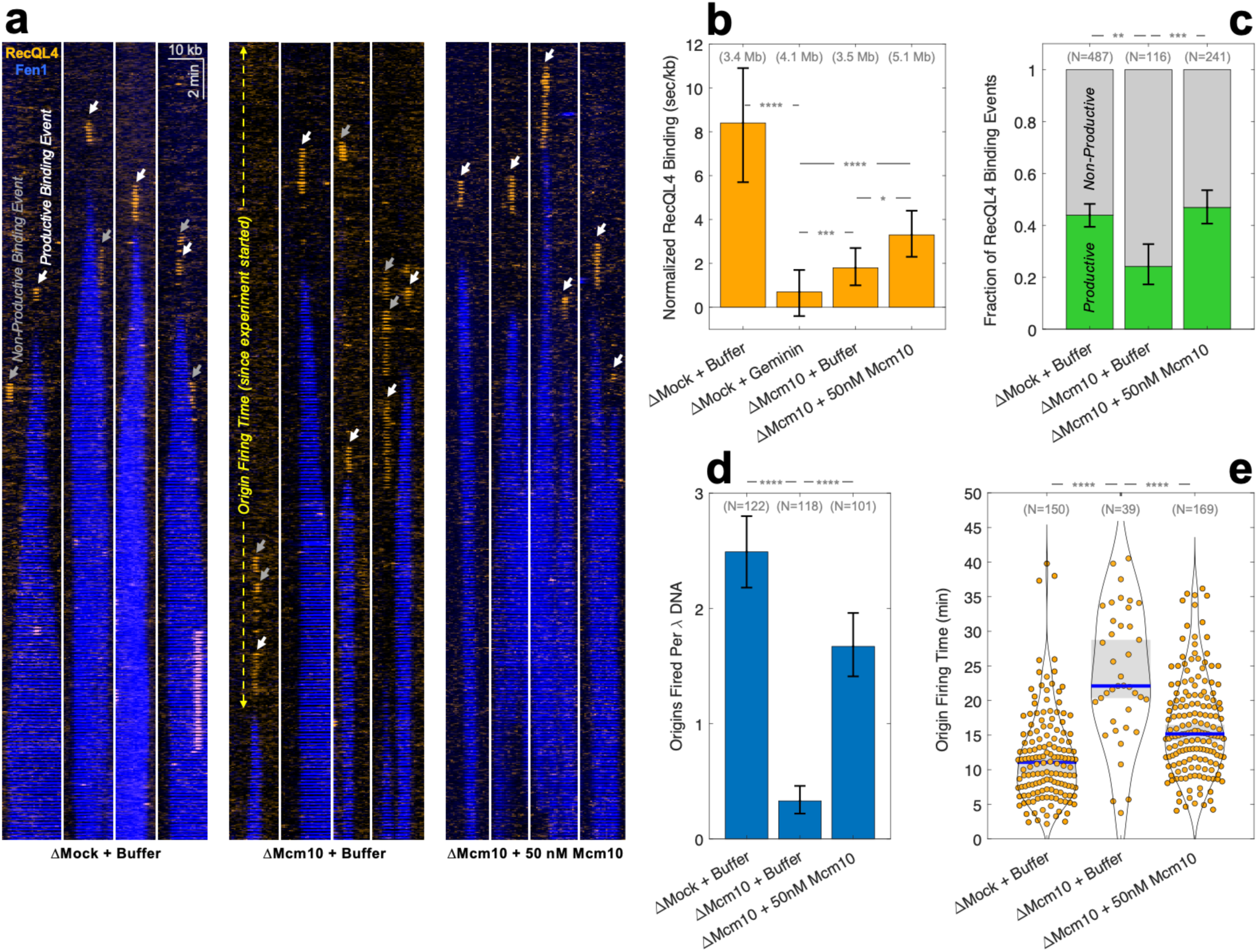
**(a)** Representative kymograms illustrating RecQL4^AF647^ (orange) binding to licensed DNA. Fen1^mKikGR^ (blue) marks nascent DNA. **(b)** Quantification of RecQL4^AF647^ binding to DNA. Total DNA length analyzed is reported in parentheses. **(c)** Fraction of productive (green) versus non-productive (gray) RecQL4^AF647^ binding events. N – number of binding events analyzed. **(d)** Number of origins fired per DNA molecule. N – number of λ DNA molecules analyzed. **(e)** Time delay between the start of the experiment (addition of nucleoplasmic extract) and origin firing. N – number of origin firing events. Blue bars and gray boxes denote the median and 95% CI. In **(b)**, **(c)**, **(d)** error-bars denote the 95% CI. Two-sample Kolmogorov-Smirnov test was used to compute p-values: ns (not significant, p > 0.05), * (p < 0.05), ** (p < 0.01), *** (p < 0.001), **** (p < 0.0001).

Interestingly, depleting Mcm10 did not completely block RecQL4 binding to chromatin (Fig. 3b), indicating that RecQL4 can be recruited to origins in a Mcm10-independent manner, albeit much less efficiently. The fraction of productive RecQL4^AF647^ binding events was significantly reduced upon depleting Mcm10 (Fig. 3c), suggesting that RecQL4 molecules recruited to DNA were less effective at promoting origin firing in the absence of Mcm10. Depleting Mcm10 did not affect the time delay between RecQL4 binding and origin firing, or the duration of non-productive RecQL4 binding events (Extended Data Fig. 3c-f).

Although 10 nM of RecQL4^AF647^ supported robust origin firing when endogenous Mcm10 was present (∼2.5 origins/DNA), depleting Mcm10 strongly supressed origin firing (∼0.3 origins/DNA) (Fig. 3d). Moreover, origin firing time was significantly delayed in Mcm10-depleted extract (∼21 minutes after the start of the experiment) compared to mock-depleted extract (∼11 min) (Fig. 3e). Both origin firing efficiency and origin firing timing were rescued by supplementing extract with 50 nM of recombinant Mcm10 (Fig. 3d-e), consistent with the idea that Mcm10 promotes efficient replication initiation by recruiting RecQL4 to origins.

### Mcm10 and RecQL4 Play Partially Redundant Roles During Replication Initiation

To dissect the division of labor between RecQL4 and Mcm10 during CMG activation, we replicated a plasmid DNA template in extract depleted of RecQL4 (ΔRecQL4), Mcm10 (ΔMcm10), or both (ΔRecQL4+ΔMcm10) (Fig. 4a). Since depleting Mcm10 or RecQL4 does not alter the speed of replication forks (Fig. 2e and Fig. 1e), any differences in replication kinetics/efficiency measured in the biochemical assay are caused by differences in origin firing. DNA replication was strongly (but not completely) suppressed in RecQL4-depleted extract (Fig. 4b lanes 5-8, compare to mock in lanes 1-4; quantification in Fig. 4d). In contrast, a stringent depletion of Mcm10 caused only a minor reduction in replication efficiency (Fig. 4b lanes 9-12, compare to lanes 1-4). Depleting both RecQL4 and Mcm10 completely abolished DNA replication (Fig. 4b lanes 13-16), consistent with the idea that both proteins are involved in CMG helicase activation. Importantly, DNA replication in double-depleted extract was efficiently rescued by near-endogenous levels (100 nM) of recombinant RecQL4 (Fig. 4b, lanes 17-20; see Fig. 4c for corresponding immunoblot), but only partially rescued by near-endogenous levels (100 nM) of recombinant Mcm10 (Fig. 4b, lanes 21-24, also see Extended Data Fig. 4a-c). These results suggest that the two proteins play distinct but partially redundant roles during CMG activation.

**Figure 4:**
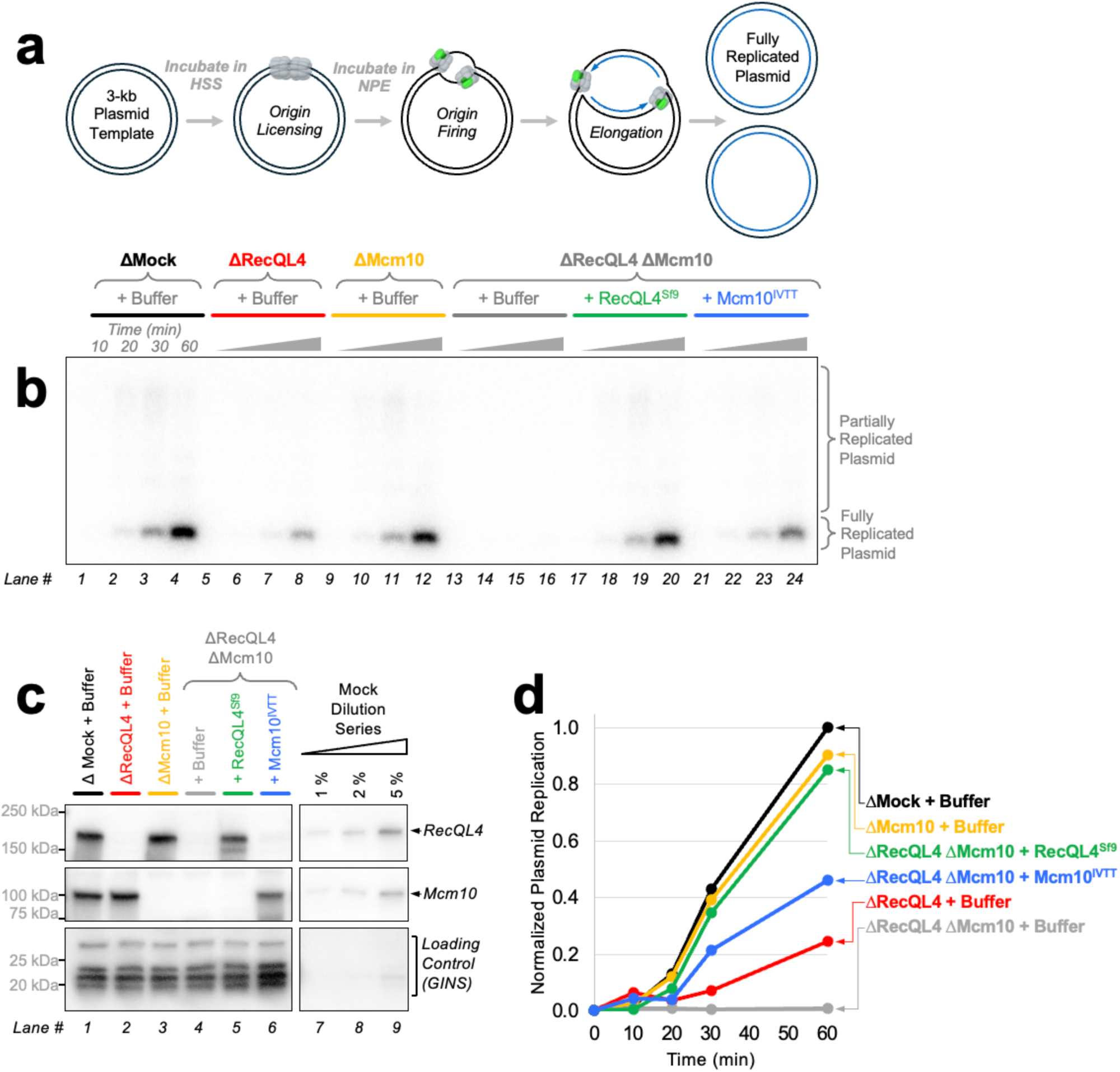
**(a)** Diagram illustrating the workflow of the ensemble plasmid replication assay (dCTP^P32^ is added to the reaction and is incorporated into nascent DNA). **(b)** Autoradiograph of agarose gel showing the results of a plasmid replication assay performed in mock-depleted, RecQL4-depleted, Mcm10-depleted, or double-depleted extract (a representative experiment of three is shown, see Extended Data Fig. 4a-c for a biological repeat). Reactions were supplemented with buffer, 100 nM of recombinant RecQL4 expressed in Sf9 insect cells (RecQL4^Sf9^), or 100 nM of Mcm10 expressed using In Vitro Transcription-Translation (RecQL4^IVTT^). Time-points were collected at 10, 20, 30, and 60 minutes. **(c)** Immunoblots of extract samples taken from reactions shown in panel **(b)**. Both Mcm10 and RecQL4 depletions are very stringent (less than 1% of endogenous protein remaining). GINS serves as a loading control. **(d)** Quantification of the nascent DNA signal for each reaction shown in panel **(b)**.

### Depleting Mcm10 Unmasks the Role of RecQL4’s Structured C-terminus in Initiation

Since RecQL4 and Mcm10 have partially redundant functions during origin firing, we hypothesized that replication initiation defects caused by RecQL4 mutations are partially masked by the presence of Mcm10. Notably, most mutations reported in RTS, BGS, and RS patients reside in RecQL4’s largely structured C-terminal portion (Fig. 5a, Extended Data Fig. 3b)^47^. To dissect the roles of RecQL4’s disordered N-terminus and structured C-terminus in initiation, we measured plasmid replication upon depleting only RecQL4 or both RecQL4 and Mcm10. Depleted extracts were supplemented with one of the following recombinant proteins: wildtype RecQL4 (RecQL4^WT^), ATPase-dead RecQL4 with a point mutation in the Walker B motif which can bind ATP but cannot hydrolyze it (RecQL4^D856A^), or a construct that contains only the disordered N-terminus (RecQL4^1-663^). Importantly, RecQL4^D856A^ mimics RTS patient mutations that impair ATPase function, while RecQL4^1-663^ mimics RTS patient mutations with a premature stop codon where the entire C-terminus of RecQL4 is missing^47^.

**Figure 5:**
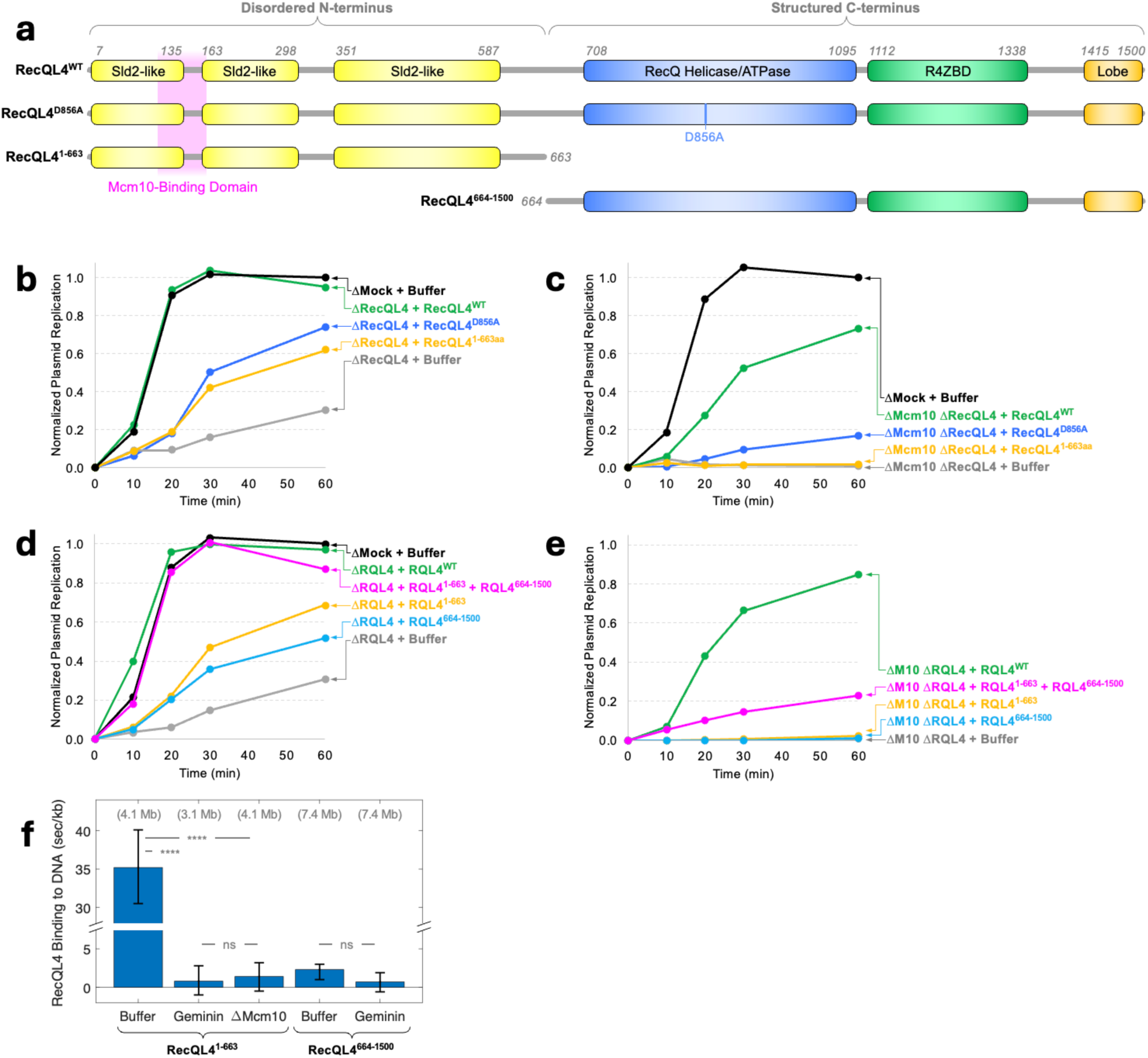
**(a)** Domain diagram for *Xenopus laevis* RecQL4 constructs used in this study. See Extended Data Fig. 1a for similar domain diagrams for wildtype human, mouse, and yeast orthologs. **(b)** Normalized signal from plasmid replication reactions performed in mock-depleted or RecQL4-depleted extract supplemented with buffer, or 100 nM of recombinant RecQL4 (wildtype in green, WalkerB mutant in blue, or N-terminal fragment in orange). A representative experiment of three is shown, see Extended Data Fig. 5a and Extended Data Fig. 5c for the raw autoradiograph and corresponding immunoblots. **(c)** Same as **(b)**, but using extract depleted of both Mcm10 and RecQL4. See Extended Data Fig. 5b for the raw autoradiograph. **(d)** Normalized signal from plasmid replication reactions performed in RecQL4-depleted extract supplemented with buffer or 100 nM of recombinant RecQL4 (wildtype in green, N-terminal fragment in orange, C-terminal fragment in blue, or both in magenta). **(e)** Same as **(d)**, but using extract depleted of both Mcm10 and RecQL4. See Extended Data Fig. 5d-f for the raw autoradiographs and immunoblots. **(f)** Quantification of RecQL4^1-663^ and RecQL4^664-1500^ binding to DNA (both protein fragments were labeled with AF647). Total DNA length analyzed is reported in parentheses. Error-bars denote the 95% CI. Two-sample Kolmogorov-Smirnov test was used to compute p-values: ns (not significant, p > 0.05), * (p < 0.05), ** (p < 0.01), *** (p < 0.001), **** (p < 0.0001).

In RecQL4-depleted extract, RecQL4^WT^ supported robust DNA replication, indistinguishable from mock (Fig. 5b, green versus black curves, also see Extended Data Fig. 5a). In the same conditions, both RecQL4^D856A^ and RecQL4^1-663^ exhibited a modest but significant delay in replication kinetics (how fast DNA is replicated), and reduced replication efficiency (how much total DNA is replicated) (Fig. 5b, blue and orange curves). These data are consistent with previous reports that the ATPase domain is not strictly required for DNA replication^53,54^.

In the absence of Mcm10, RecQL4^D856A^ exhibited much lower replication efficiency than wildtype (Fig. 5c, blue versus green curves, also see Extended Data Fig. 5b-c). This indicates that Mcm10 largely compensates for the defect caused by inactivating RecQL4’s ATPase. Surprisingly, RecQL4^1-663^ was incapable of supporting replication in Mcm10-depleted extract (Fig. 5c, orange curve). This suggests that RecQL4’s C-terminus encodes for another function (distinct from the ATPase) which becomes critical for initiation in the absence of Mcm10. We conclude that Mcm10 masks replication defects caused by RecQL4 mutants which mimic RTS patient mutations.

### RecQL4’s N-terminus and C-terminus Both Synergize with Mcm10 to Promote Origin Firing

To better understand the molecular basis of RecQL4 syndromes, we sought to dissect the function of RecQL4’s N and C terminal domains. To account for Mcm10’s role in initiation, we measured plasmid replication in RecQL4-depleted or double-depleted extract supplemented with buffer, RecQL4^1-663^, RecQL4^664-1500^, or RecQL4^WT^.

RecQL4^1-663^ supported some DNA replication in RecQL4-depleted extract, but not in double-depleted extract (Fig. 5d-e, orange curve; also see Extended Data Fig. 5d-f). To explain this, we hypothesized that Mcm10 recruits RecQL4^1-663^ to origins via the known RecQL4-Mcm10 interaction (Fig. 5a, magenta). Consistent with this idea, depleting Mcm10 abolished RecQL4^1-663^ binding to DNA (Fig. 5f).

Similarly, RecQL4^664-1500^ supported some replication in RecQL4-depleted extract, but not in double-depleted extract (Fig. 5d-e, blue curve; also see Extended Data Fig. 5d-e). Importantly, the ability of RecQL4^664-1500^ to support replication entirely depends on its ATPase activity (Extended Data Fig. 5g, blue versus red curves). These observations cannot be explained by a model where Mcm10 recruits RecQL4^664-1500^ to origins, because Mcm10 interacts with the N-terminus of RecQL4 and RecQL4^664-1500^ binding to DNA is already very low even without depleting Mcm10 (Fig. 5f).

Taken together, these data indicate that the N and C terminal domains of RecQL4 separately synergize with Mcm10 to promote replication initiation. In the case of RecQL4’s C-terminus, this synergy depends on its ATPase activity.

### RecQL4’s N-terminus and C-terminus Synergize with Each Other During Initiation

Although neither RecQL4^1-663^ nor RecQL4^664-1500^ supported origin firing in double-depleted extract (Fig. 5e, orange and blue curves), when added together they supported some DNA replication (Fig. 5e, magenta curve). This synergy is even more pronounced in the presence of Mcm10, when RecQL4^1-663^ and RecQL4^664-1500^ supported replication as well as RecQL4^WT^ (Fig. 5d, magenta versus green curves). Taken together, these data suggest that (i) RecQL4’s N-terminus plays a non-catalytic role in initiation, which is not sufficient for origin firing in the absence of Mcm10; (ii) RecQL4’s C-terminus plays a catalytic role in initiation, which is not sufficient for origin firing in the absence of Mcm10; and (iii) RecQL4’s N-terminus and C-terminus synergize to promote detectable levels of replication even in the absence of Mcm10.

### RecQL4’s C-terminus Mediates a Previously Unknown Interaction with Mcm7

The dramatic difference in replication efficiency between RecQL4^D856A^ and RecQL4^1-663^ in Mcm10-depleted extract (Fig. 5c, blue versus orange curves) suggests that RecQL4’s C-terminus encodes a function distinct from ATP hydrolysis and DNA binding. AlphaFold3 predicts that a previously unannotated portion of RecQL4’s C-terminus should interact with the winged-helix domain of Mcm7 (Fig. 6a-b, also see Extended Data Fig. 6a-b for human/mouse models). We hypothesized that this interaction is responsible for the Mcm10-independent recruitment of RecQL4^664-1500^ to the pre-activation complex.

**Figure 6:**
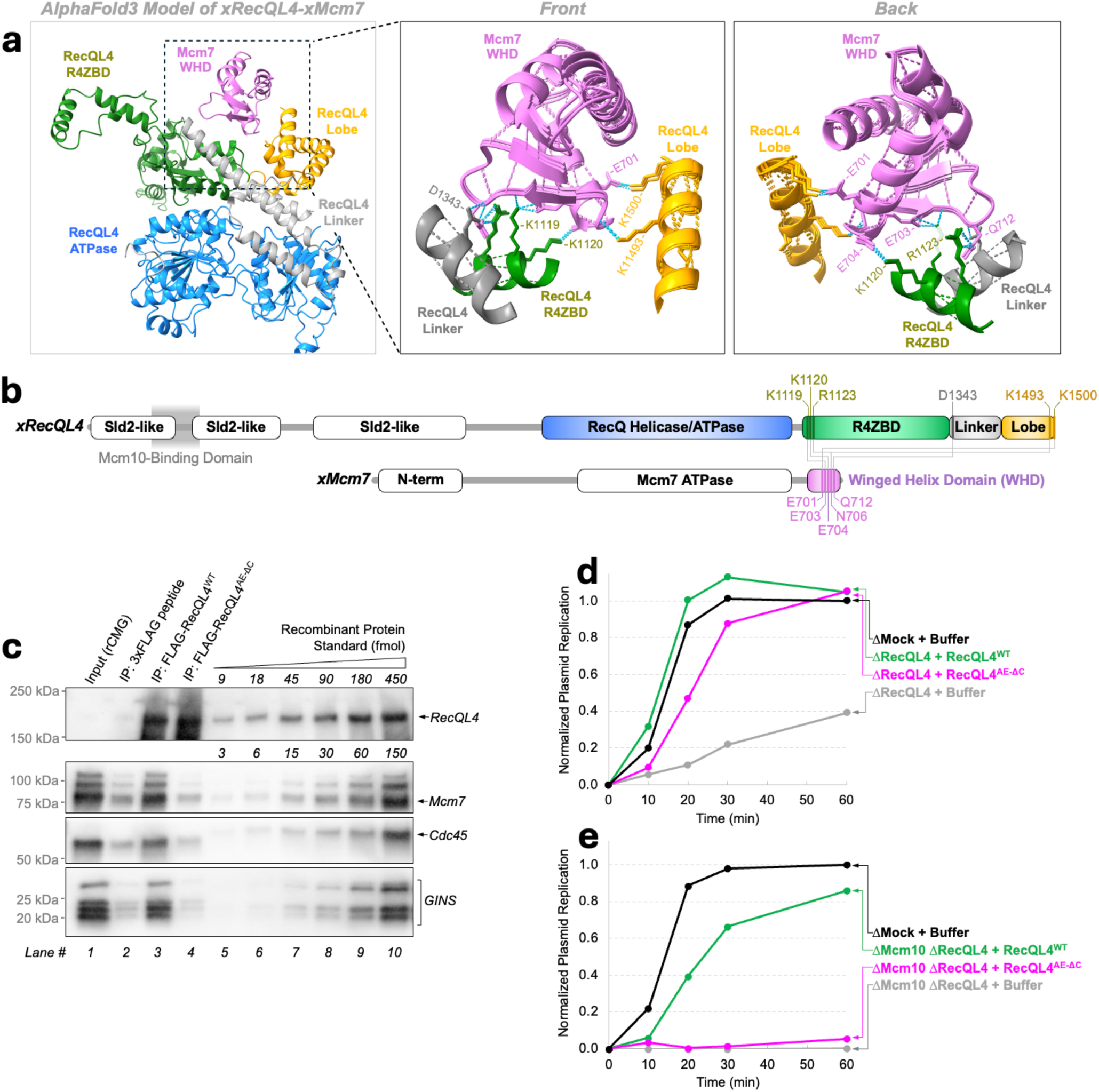
**(a)** Left: AlphaFold3-predicted structure of frog Mcm7’s winged-helix-domain (magenta) bound to the structured portion of RecQL4. Extended Data Fig. 6a shows similar predictions for human and mouse orthologs. Middle/Right: magnified view of the top three predicted structures (overlayed) highlighting key interactions. For clarity, some parts of the structure are not shown. **(b)** Diagram of frog RecQL4 and Mcm7 highlighting residues involved in AlphaFold3-predicted interactions, using the same color scheme as in panel **(a)**. **(c)** Immunoblot showing the results of the co-immunoprecipitation experiment between recombinant frog CMG (untagged) and recombinant FLAG-tagged RecQL4 (wildtype, or RecQL4^AE-ΔC^ - a mutant predicted to no longer interact with Mcm7). **(d)** Normalized signal from plasmid replication reactions performed in RecQL4-depleted extract supplemented with buffer or 100 nM of recombinant RecQL4^WT^ (green) or RecQL4^AE-ΔC^ (magenta). **(e)** Same as **(d)**, but using extract depleted of both Mcm10 and RecQL4. See Extended Data Fig. 6c-e for raw autoradiographs and immunoblots corresponding to panels **(d)** and **(e)**.

As reported previously, truncating the last 92 amino-acids from the human RecQL4 impairs its ability to bind ssDNA and unwind dsDNA *in vitro*^55^. Therefore, truncating the extreme C-terminus of RecQL4 is not a suitable way of specifically ablating the interaction with Mcm7. Guided by the AlphaFold3 prediction, we sought to disrupt the RecQL4-Mcm7 interaction by introducing two point-mutations (K1119A, K1120E) and deleting the last 9 amino-acids of RecQL4. We refer to this Mcm7-interaction deficient mutant as RecQL4^AE-ΔC^. *In vitro*, recombinant CMG co-immunoprecipitated with purified RecQL4^WT^ but not with RecQL4^AE-ΔC^ (Fig. 6c), indicating that the RecQL4-Mcm7 was effectively disrupted in this mutant.

In the presence of Mcm10, RecQL4^AE-ΔC^ supported replication nearly as well as wildtype (Fig. 6d magenta curve, also see Extended Data Fig. 6c), indicating that disrupting the Mcm7-RecQL4 interaction does not impair RecQL4’s ability to activate CMG. However, in the absence of Mcm10, RecQL4^AE-ΔC^ supported a negligible amount of replication (Fig. 6e, magenta curve, also see Extended Data Fig. 6d-e), consistent with the idea that the interaction with Mcm7 serves as an alternative way of recruiting RecQL4 to the pre-activation complex.

## DISCUSSION

### Proposed Model of CMG Helicase Activation in Metazoa

Based on the data presented here, we propose a new model of CMG activation in metazoa (Fig. 7). Fig. 7a-b depicts a model of the pre-activation complex containing two inactive CMGs encircling dsDNA, and a dimer of DONSON that “staples” them. We found that Mcm10 is recruited to the pre-activation complex first, as shown in Fig. 7c. AlphaFold3 predicts that the well-folded core of the *Xenopus* Mcm10 (containing a ssDNA binding domain^56^) should bind to the N-terminus of Mcm4. The placement of Mcm10 agrees with a recent yeast CMG-Mcm10 structure^19^, and is consistent with cross-linking mass spectrometry data^57^. Our data indicate that Mcm10 recruits RecQL4 to the pre-activation complex (Fig. 7d), likely via the known interaction between Mcm10 and RecQL4’s N-terminus^35^.

**Figure 7:**
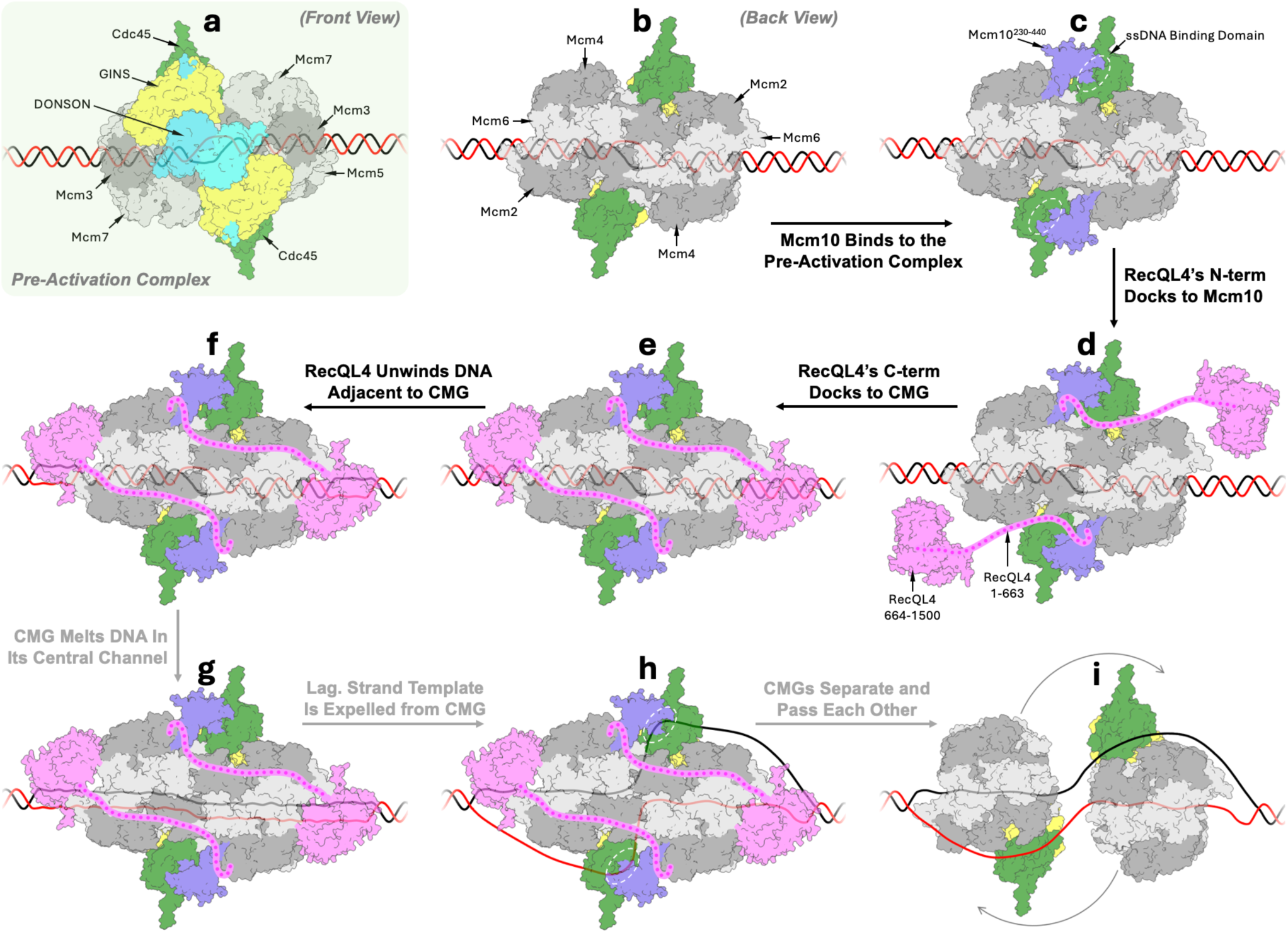
Proposed model of CMG activation in metazoa. Thick black arrows highlight events supported by our data. Thich gray arrows indicate events previously known to occur during CMG activation. **(a)** Front view of the pre-activation complex consisting of two head-to-head CMGs on dsDNA bound by a DONSON dimer (light blue). Model was obtained by aligning a human CMG structure (PDB 6XTX) to a structure of frog (CMG•DONSON)_2_ which lacks the Mcm2-7 ATPase domains (PDB 8Q6O). CMG central channels of CMGs are offset in the PDB 8Q6O structure, suggesting that the DNA inside the pre-activation complex should be partially melted. **(b)** Back view of the pre-activation complex model (DONSON and GINS are not visible here). **(c)** AlphaFold3-predicted docking of Mcm10’s OB-fold domain known to bind ssDNA (white oval). AlphaFold3 did not consistently predict the docking of other Mcm10 domains. **(d)** The disordered N-terminus of RecQL4 (magenta) docks to Mcm10 (blue). **(e)** The structured C-terminus of RecQL4 (magenta) docks to CMG (AlphaFold3 prediction). **(f)** RecQL4’s melts dsDNA at the periphery of the pre-activation complex. **(g)** We hypothesize that Mcm10 and RecQL4’s N-terminus allosterically stimulate CMG’s ATPase/helicase activity. We hypothesize that the two CMGs in a head-to-head orientation exert force/torque on dsDNA inside their central channels, which ultimately leads to the melting of origin DNA (similar to a model proposed by the O’Donnell Lab for yeast CMG activation^21^). **(h)** The lagging-strand template is expelled from CMG’s central channel. **(i)** The two CMGs separate from each other and begin translocating along different template DNA strands. Mcm10 and/or RecQL4 may briefly bind the N-terminal face of Mcm2-7 to prevent the re-dimerization of the two CMGs.

AlphaFold3 consistently docks RecQL4’s structured C-terminus (660-1500 a.a.) to CMG’s C-terminal side, in proximity to dsDNA (Fig. 7e). Since RecQL4 can unwind short DNA oligos *in vitro*^58^, we propose that RecQL4 promotes CMG activation by melting dsDNA on either side of the pre-activation complex (Fig. 7f). This helps explain why RecQL4’s ATPase activity is important for CMG helicase activation. In the absence of Mcm10, RecQL4-Mcm7 interaction serves as a backup means of recruiting RecQL4 to origins.

We found that RecQL4’s N-terminus and Mcm10 both play non-catalytic roles in CMG activation, and both synergize with the catalytic function of RecQL4’s C-terminus. Yeast Mcm10 was shown to allosterically stimulate CMG’s ATPase-helicase activity *in vitro*^21,59^. Our data is consistent with a model where both Mcm10 and RecQL4^1-663^ stimulate CMG’s ATPase-helicase activity, thus promoting DNA melting inside CMGs central channel (Fig. 7g).

During CMG activation, the lagging strand template must be expelled from CMG’s central channel. We speculate that Mcm10’s ssDNA binding domain captures the excluded strand and stabilizes it long enough to facilitate the activation of both CMG complexes (Fig. 7h, white ovals). Finally, the two CMGs separate from each other, and this is accompanied by Mcm10, RecQL4, and DONSON dissociation. As proposed previously^18,21^, the head-to-head CMG orientation ensures that each origin fires only when both CMGs are fully activated and can pass each other unobstructed on different DNA strands (Fig. 7i).

### Division of Labor Between RecQL4 and Mcm10 During Replication Initiation

Although both RecQL4 and Mcm10 were previously implicated in metazoan replication initiation, the exact roles of these proteins have long been debated. Recently, we showed that RecQL4 participates in CMG helicase activation^17^. Here, we show that Mcm10 also participates in CMG helicase activation, and that it synergizes with RecQL4. Our data indicate that Mcm10 and RecQL4 are partially redundant during CMG activation, explaining why RecQL4-depleted *Xenopus* extracts still support significant levels of DNA replication^17^, and why some RecQL4-knockout cells maintain viability in cell culture^60^.

Why do metazoa employ two distinct initiation factors to drive CMG activation? It is possible that Mcm10 and RecQL4 may act on different classes of origins. For example, either protein alone may be sufficient to promote CMG activation at origins in euchromatin, whereas firing origins in heterochromatin may require both proteins acting in concert. Alternatively, Mcm10 or RecQL4 may be selectively recruited to specific loci. For example, RecQL4 can bind DNA secondary structures such as G4 quadruplexes, and this may regulate replication timing^61^. Future studies mapping Mcm10-dependent versus RecQL4-dependent origins will further elucidate the division of labor between these two CMG activators.

### Mechanistic Differences and Similarities Between Yeast and Metazoa

To date, the mechanism of CMG helicase activation is best understood in yeast. It is widely accepted that yeast Mcm10 plays a central role in CMG activation^18^, whereas Sld2 (the yeast homolog of RecQL4) is thought to participate in CMG assembly^24^. Our data reveal that unlike yeast, metazoan CMG activation involves the concerted action of both Mcm10 and RecQL4. Importantly, we show that RecQL4’s structured C-terminus is critical for efficient origin firing, especially in the absence of Mcm10. This fragment of RecQL4 is unique to metazoa and includes the ATPase domain, the DNA-binding domain (R4ZBD), and the Mcm7-interaction motif we identified (Fig. 6b, also see Extended Data Fig. 1A, Extended Data Fig. 7).

The role of Mcm10 and RecQL4 in metazoan replication initiation has been highly debated because these proteins are involved in several other aspects of genome maintenance^27,28,62^. In hindsight, previous conflicting reports about the roles of RecQL4/Mcm10 can be explained by the fact that Mcm10 and RecQL4 are partially redundant during CMG activation, as shown here.

### Mcm10 Does Not Travel with the Replication Fork During Unperturbed Replication

Sometimes called “the most mysterious replication protein”, Mcm10 has been implicated in numerous aspects of DNA replication, ranging from initiation, elongation^62^, termination^63^, replication fork reversal ^64^, and replication fork restart^27,65^. These functions imply that Mcm10 travels with the replication fork. It is widely accepted that Mcm10 is a constitutive replisome subunit, with supporting evidence coming from reports of Mcm10 enrichment at/near sites of active replication^62^. However, there are also studies reporting little to no enrichment of Mcm10 at replication forks^23^. A recent single-molecule study with purified yeast proteins, found that Mcm10 travels with CMG, but those experiments did not recapitulate CMG loading/activation and did not contain fully assembled replisomes^66^. So far, there have been no studies that directly visualized Mcm10 at active replication forks.

Our data indicates that Mcm10 is recruited to the pre-activation complex and that Mcm10 dissociates from the origin shortly before the start of DNA synthesis. In >90% of cases we do not observe Mcm10 travel with active replication forks (Fig. 1). In the absence of any additional perturbations, we rarely observe Mcm10 transiently associating with active replisomes. We propose that following CMG activation, Mcm10 is displaced from its binding site at the front of CMG by other replisome subunits. For example, Claspin and Timeless-TIPIN also bind at the front of the CMG complex and could prevent Mcm10 re-binding^67^. It is also possible that Mcm10 does not associate with replisomes during unperturbed DNA replication but is conditionally recruited to CMG during replication stress. Future studies are needed to explore this idea.

### Replication Initiation Defects Caused by Pathological RecQL4 Mutations

The field lacks a clear understanding of how mutations in RecQL4 lead to the phenotypes reported in patients with Rothmund-Thomson, Baller-Gerold, and RAPADILINO syndromes. Our data is consistent with the idea that RecQL4 disease phenotypes may be attributed to its role in replication initiation.

A large fraction of disease-associated RecQL4 mutants contain premature stop codons or large deletions predicted to result in low mRNA/protein levels^68^. Similarly, many point mutations or small deletions may also destabilize RecQL4 folding/stability. For such hypomorphic mutations, the exact location of the mutation (Sld2-like domain, ATPase domain, R4ZBD, etc.) should be interpreted with caution. The one exception are premature stop codons in the last exon, which are usually not subject to nonsense-mediated decay^69^. There are two such mutations reported in Rothmund-Thompson Syndrome patients: p.Gln1175*^70^ and p.Thr1200Arg*fs*\*26^47^. They are expected to produce almost full length RecQL4 at nearly-normal levels, yet are pathological^68^. Here we report that the unannotated extreme C-terminus of RecQL4 mediates the interaction with Mcm7 (Fig. 6). We propose that this interaction docks RecQL4’s ATPase-helicase to the rear of CMG, proximal to dsDNA (Fig. 7d-e).

Our data show that all domains of RecQL4 play a role in replication initiation, therefore any patient-derived mutations that impair protein function or stability are expected to reduce origin firing efficiency. Since Mcm10 can partially compensate for the loss of RecQL4’s ATPase activity/domain, we speculate that Mcm10 may be overexpressed in some patient tissues to compensate for pathologic RecQL4 mutations.

### Limitations

All experiments were repeated at least twice with different extract preparations, a representative experiment is shown in the figures. Different extract preparations may exhibit some variations in DNA replication kinetics which could be caused by small changes in firing factor concentrations.

According to the OpenCell database^71^, the cellular concentrations of Mcm10 (∼50 nM) and RecQL4 (∼50 nM) are lower than in *Xenopus* egg extract (∼250 nM for RecQL4 and ∼100 nM for Mcm10 in our biochemical reactions). Plasmid replication reactions were performed with 50-100 nM of unlabeled recombinant Mcm10 or RecQL4, which was saturating for the ensemble assays.

Single-molecule experiments were performed at 10 nM of Mcm10^AF647^ or RecQL4^AF647^ to maintain an acceptable signal-to-noise ratio. Although this concentration is lower than that of endogenous proteins, these experiments provided a sensitive way of quantifying the relative binding affinity for different mutants under different depletion conditions.

In single-molecule experiments conducted with 10 nM of RecQL4^AF647^, origin firing efficiency was very sensitive to Mcm10 concentration (Fig. 3d). This is consistent with the idea that Mcm10-independent recruitment of RecQL4 to origins is rate-limiting at low RecQL4 concentrations. In ensemble replication experiments conducted with either endogenous RecQL4 (∼250 nM) or recombinant RecQL4 (100 nM), depleting Mcm10 caused only a mild defect in origin firing (Fig. 4b, Extended Data Fig. 4a). This is consistent with the idea that Mcm10-independent recruitment of RecQL4 to origins is not rate-limiting at high RecQL4 concentrations.

The RecQL4-Mcm7 interaction is relatively weak, as stringent washes decrease the efficiency of co-precipitation *in vitro* (data not shown). This is consistent with our measurement of very low recruitment of RecQL4^664-1500^ to origins in Mcm10-depleted extract (Fig. 5f).

## Acknowledgments

We thank Karlene Cimprich, Joanna Wysocka, Nicole Martinez, Kate MacDonald, Ulrike Kuehbacher, Dhruva Deshpande, and Benjamin Duewell for suggestions based on their critical reading of the manuscript. We thank Scott Berger for feedback on the manuscript and for assistance in raising RecQL4 antibodies. We thank Justin Sparks for the gift of a plasmid encoding the full-length frog Mcm10. R.T. was supported in part by the Stanford School of Medicine Dean’s fellowship. L.A.S. was supported in part by a Blavatnik Family Fellowship Fund. G.C. is supported by an NSF CAREER Award (2144481), an NIGMS R35 award (GM147060), and an American Cancer Society seed grant (228425).

## Author Contributions

G.C., wrote the manuscript with input from R.T.; R.T. and G.C. conceived the project; R.T. carried out all ensemble and single-molecule experiments; L.A.S. raised/validated RecQL4 custom antibodies, optimized the Mcm10 immunodepletion protocol, and cloned Mcm10 for expression via IVTT; D.S. raised/validated the Mcm10 and TopBP1 custom antibodies; G.C. wrote MATLAB code to analyze single-molecule experiments; G.C. and R.T. analyzed single-molecule data. G.C. performed bioinformatics analysis.

---

**Extended Data Figure 1.**
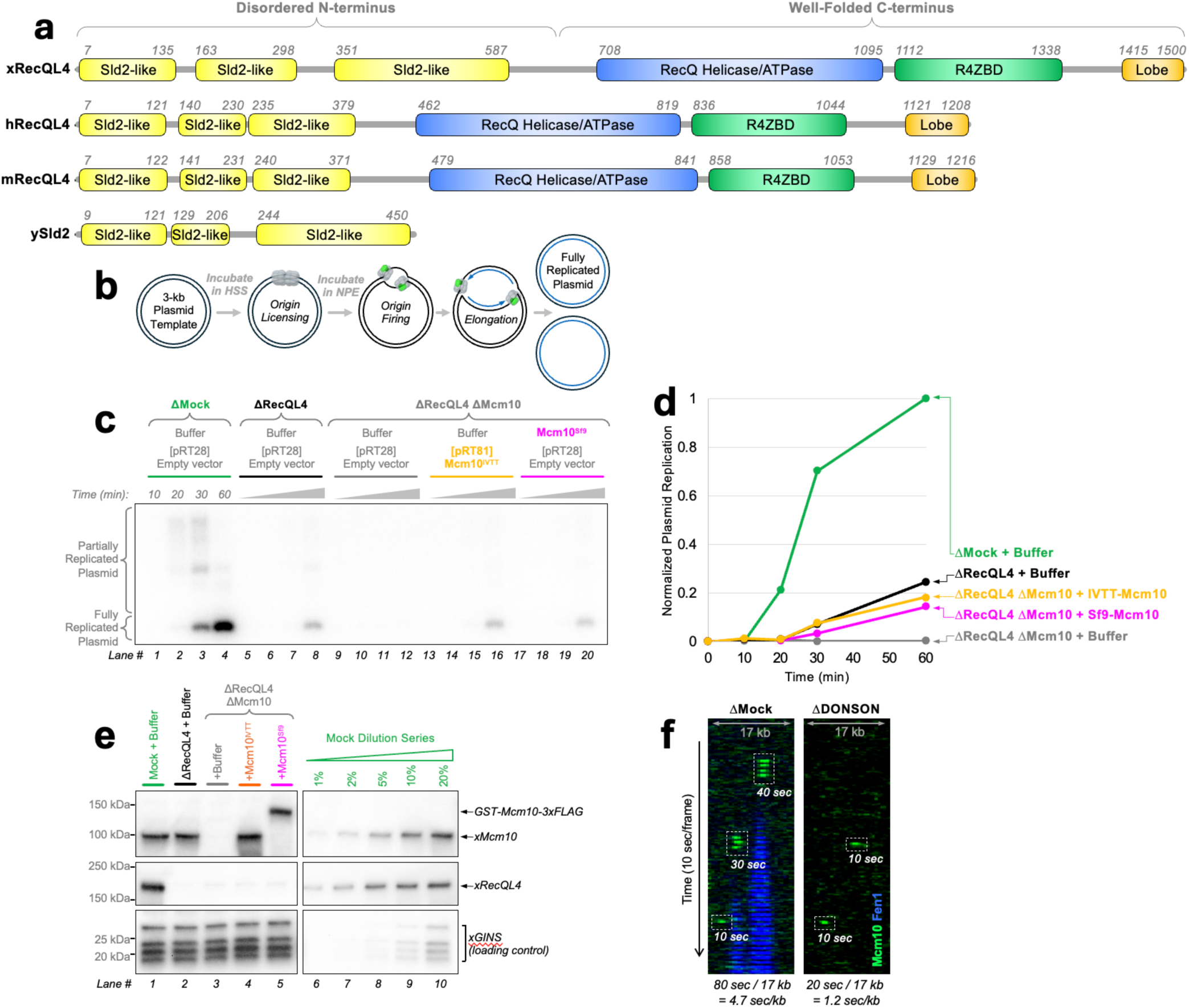
**(a)** Domain diagram for frog (x), human (h), and mouse (m) RecQL4 compared to yeast (y) Sld2. ATPase (blue) and R4ZBD (green) domain boundaries for human RecQL4 are taken from a structural study the Kisker lab^1^ and extrapolated to its orthologs using multiple-sequence alignment (see Extended Data Fig. 7). Sld2-like region boundaries (yellow) are defined here based on the multiple-sequence alignment and serve as a rough indicator of sequence conservation. C-terminal lobe boundaries (orange) were defined based on AlphaFold3 predictions and multiple-sequence alignment (see Extended Data Fig. 6a-b). **(b)** Diagram illustrating the workflow of the ensemble plasmid replication assay (dCTP^P32^ is added to the reaction and is incorporated into nascent DNA). **(c)** Autoradiograph of agarose gel showing the results of a plasmid replication assay performed in mock-depleted, RecQL4-depleted, or Mcm10-and-RecQL4-depleted extract. Reactions were supplemented with buffer, 100 nM of recombinant Mcm10 expressed in insect cells (Mcm10^Sf9^), or 100 nM of Mcm10 expressed using In Vitro Transcription-Translation (Mcm10^IVTT^). Time-points were collected at 10, 20, 30, and 60 minutes. **(d)** Quantification of the nascent DNA signal for each reaction shown in panel **(c)**. **(e)** Immunoblots of extract samples taken from reactions shown in panel **(c)**. Both Mcm10 and RecQL4 depletions are very stringent (less than 1% of endogenous protein remaining). GINS serves as a loading control. Double-depleted extract does not support any DNA replication. Both Mcm10^Sf9^ and Mcm10^IVTT^ support similar levels of DNA replication as RecQL4-depleted extract, indicating that the recombinant protein preparations have similar biochemical activity as the endogenous Mcm10. **(f)** Representative kymograms illustrating Mcm10^AF647^ binding to licensed DNA in mock-depleted or DONSON-depleted extract. To calculate the Mcm10 binding time on DNA, the durations of all binding events are added, and normalized by the length of the DNA molecule. This is done for several stretched λ DNA molecules (typically 100-200 DNA molecules). This measurement is corrected for non-specific binding of Mcm10^AF647^ off the DNA by performing a similar analysis in regions of interest that contain no DNA molecules, as previously described^2^.

**Extended Data Figure 2.**
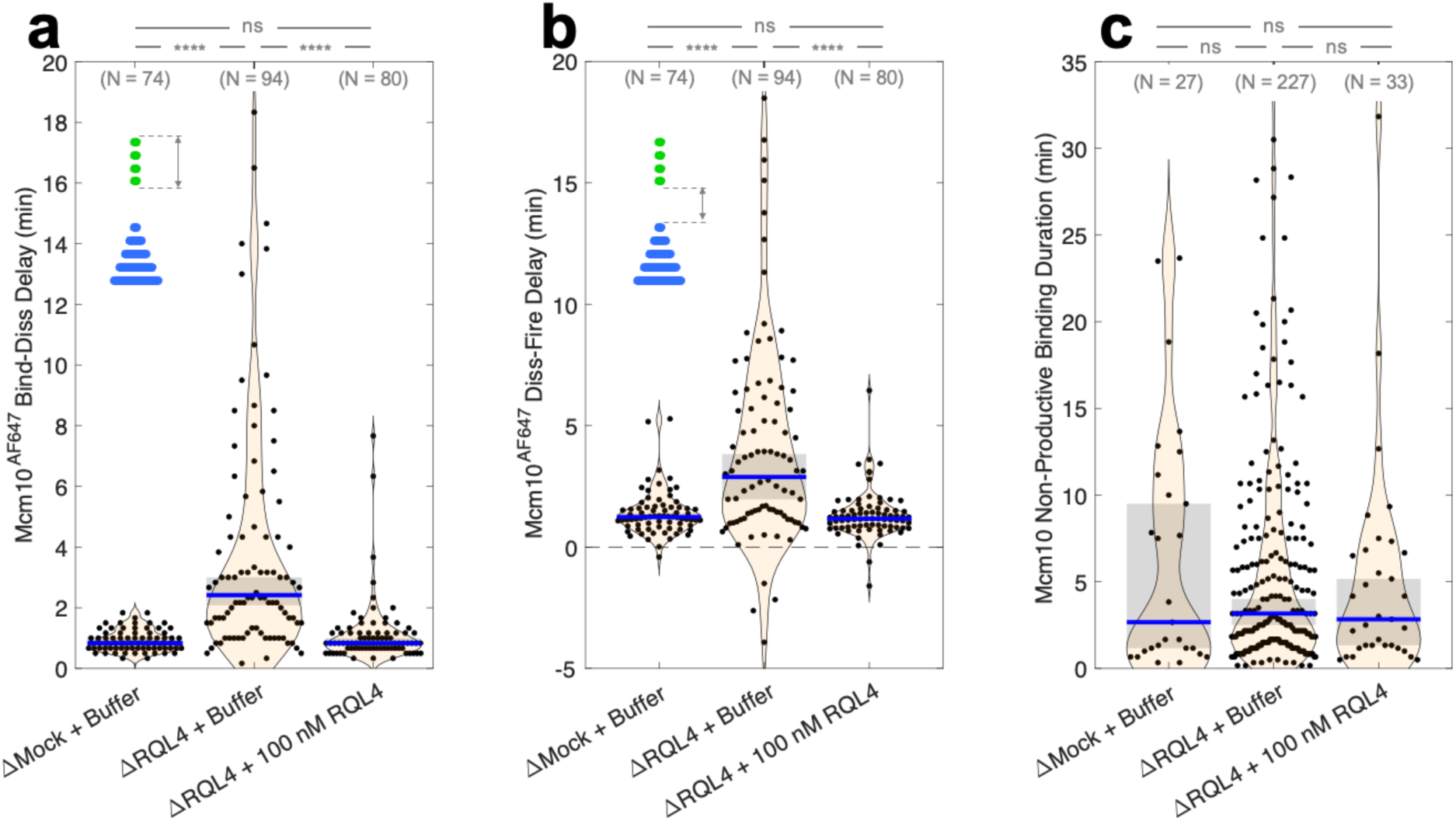
**(a)** Distribution of time delays between Mcm10^AF647^ binding and dissociation for productive binding events (i.e. the duration of productive binding events). **(b)** Distribution of time delays between Mcm10^AF647^ dissociation and origin firing. **(c)** Distribution of time delays between Mcm10^AF647^ binding and dissociation for non-productive binding events (i.e. the duration of non-productive binding events). In all panels, N denotes number of binding events analyzed; blue bars and gray boxes denote the median and 95% confidence interval for the median estimated via bootstrapping; two-sample Kolmogorov-Smirnov test was used to compute p-values: ns (not significant, p > 0.05), * (p < 0.05), ** (p < 0.01), *** (p < 0.001), **** (p < 0.0001).

**Extended Data Figure 3.**
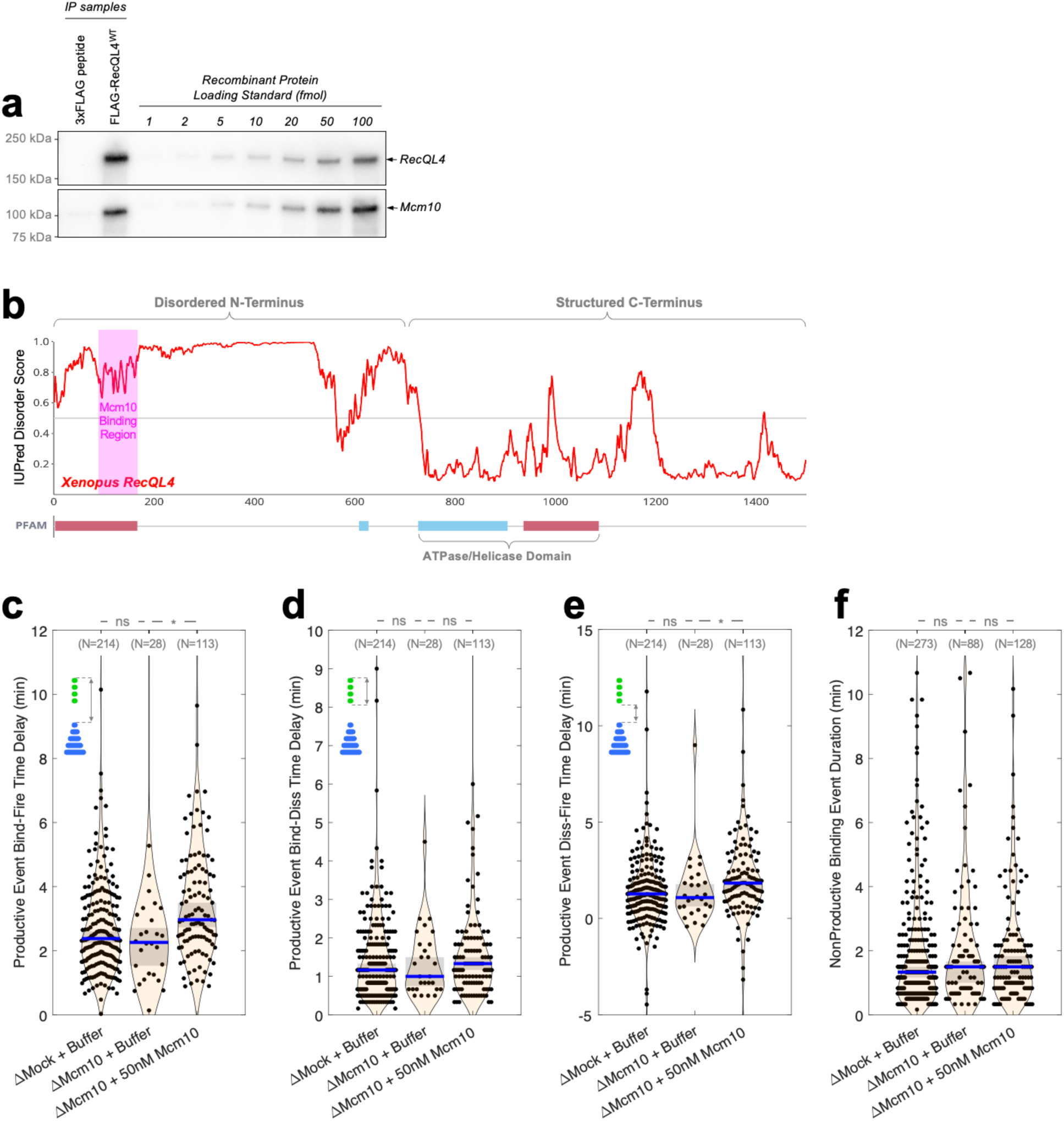
**(a)** Immunoblot showing the results of the co-immunoprecipitation experiment between recombinant frog Mcm10 (untagged) and recombinant FLAG-tagged RecQL4. 3xFLAG peptide was used as a negative control. A serial titration of both proteins was performed to estimate co-IP efficiency. **(b)** Predicted disorder score for frog RecQL4 calculated using the AIUPred server^3^. The Mcm10 binding region identified previously^4^ is shown in magenta. **(c)** Distribution of time delays between RecQL4^AF647^ binding and origin firing for productive binding events. **(d)** Distribution of time delays between RecQL4^AF647^ binding and dissociation for productive binding events (i.e. the duration of productive binding events). **(e)** Distribution of time delays between RecQL4^AF647^ dissociation and origin firing. **(f)** Distribution of time delays between RecQL4^AF647^ binding and dissociation for non-productive binding events (i.e. the duration of non-productive binding events). In panels **(c)** through **(f)**, N denotes number of binding events analyzed; blue bars and gray boxes denote the median and 95% confidence interval for the median estimated via bootstrapping; two-sample Kolmogorov-Smirnov test was used to compute p-values: ns (not significant, p > 0.05), * (p < 0.05), ** (p < 0.01), *** (p < 0.001), **** (p < 0.0001).

**Extended Data Figure 4.**
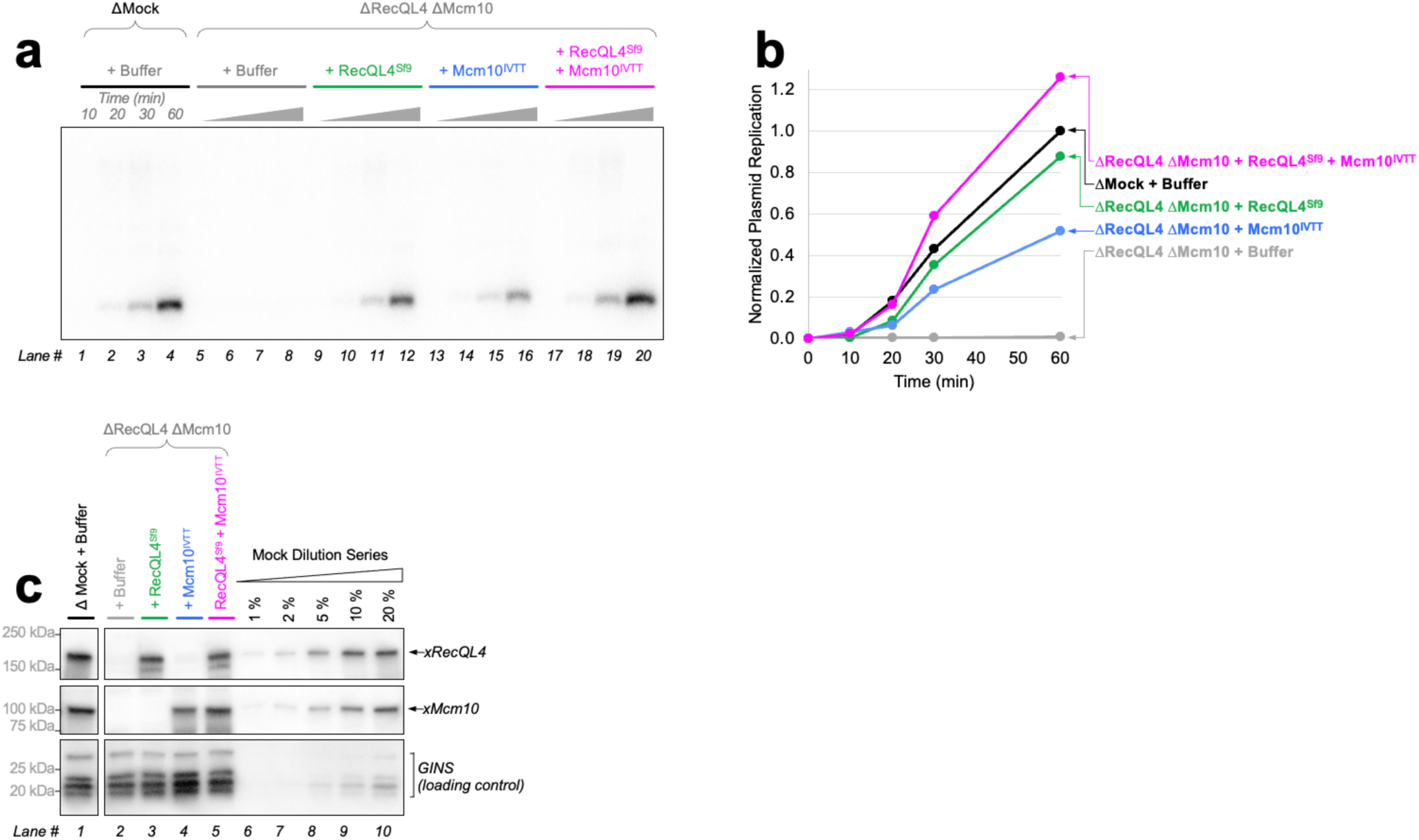
Replicate of the experiment shown in Fig. 4, focusing on the double-depletion condition. **(a)** Autoradiograph of agarose gel showing the results of a plasmid replication assay performed in mock-depleted, or Mcm10-and-RecQL4-depleted extract. Reactions were supplemented with buffer, 100 nM of recombinant RecQL4 expressed in Sf9 insect cells (RecQL4^Sf9^), 100 nM of Mcm10 expressed using In Vitro Transcription-Translation (RecQL4^IVTT^), or both RecQL4^Sf9^ and RecQL4^IVTT^. **(b)** Quantification of the nascent DNA signal for each reaction shown in panel **(a)**. **(c)** Immunoblots of extract samples taken from reactions shown in panel **(a)**. GINS serves as a loading control.

**Extended Data Figure 5.**
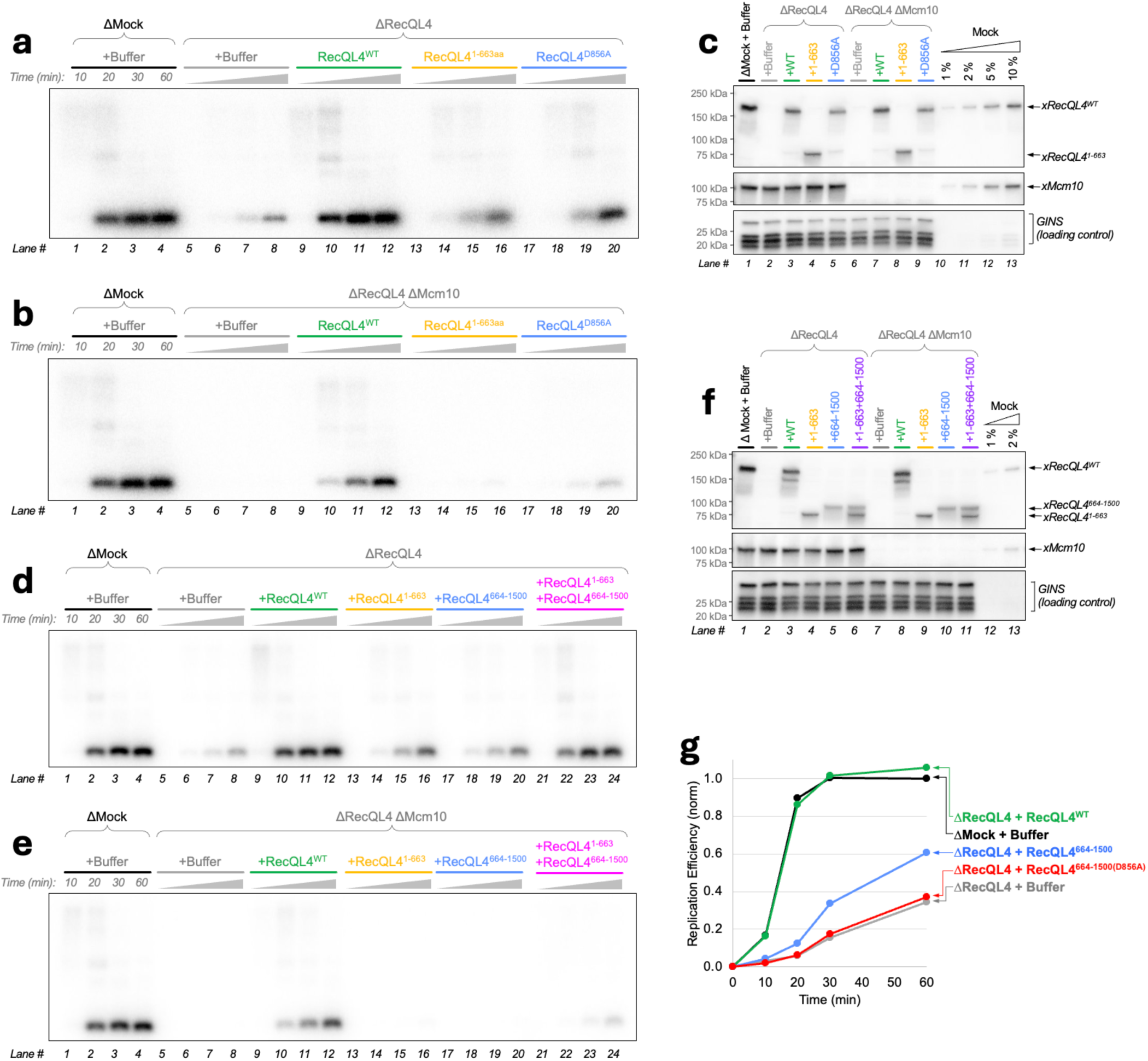
**(a)** Raw autoradiograph corresponding to data shown in Fig. 5b. **(b)** Raw autoradiograph corresponding to data shown in Fig. 5c. **(c)** Immunoblots of extract samples taken from reactions shown in panels **(a)** and **(b)**. GINS serves as a loading control. **(d)** Raw autoradiograph corresponding to data shown in Fig. 5d. **(e)** Raw autoradiograph corresponding to data shown in Fig. 5e. **(f)** Immunoblots of extract samples taken from reactions shown in panels **(d)** and **(e)**. GINS serves as a loading control. **(g)** Normalized signal from plasmid replication reactions performed in mock-depleted or RecQL4-depleted extract supplemented with buffer, 100 nM of full length RecQL4 (RecQL4^WT^, green), 100 nM of the structured C-terminal domain (RecQL4^664-1500^, blue), or 100 nM of the C-terminal domain with a mutated Walker B motif (RecQL4^664-1500(D856A)^, red).

**Extended Data Figure 6.**
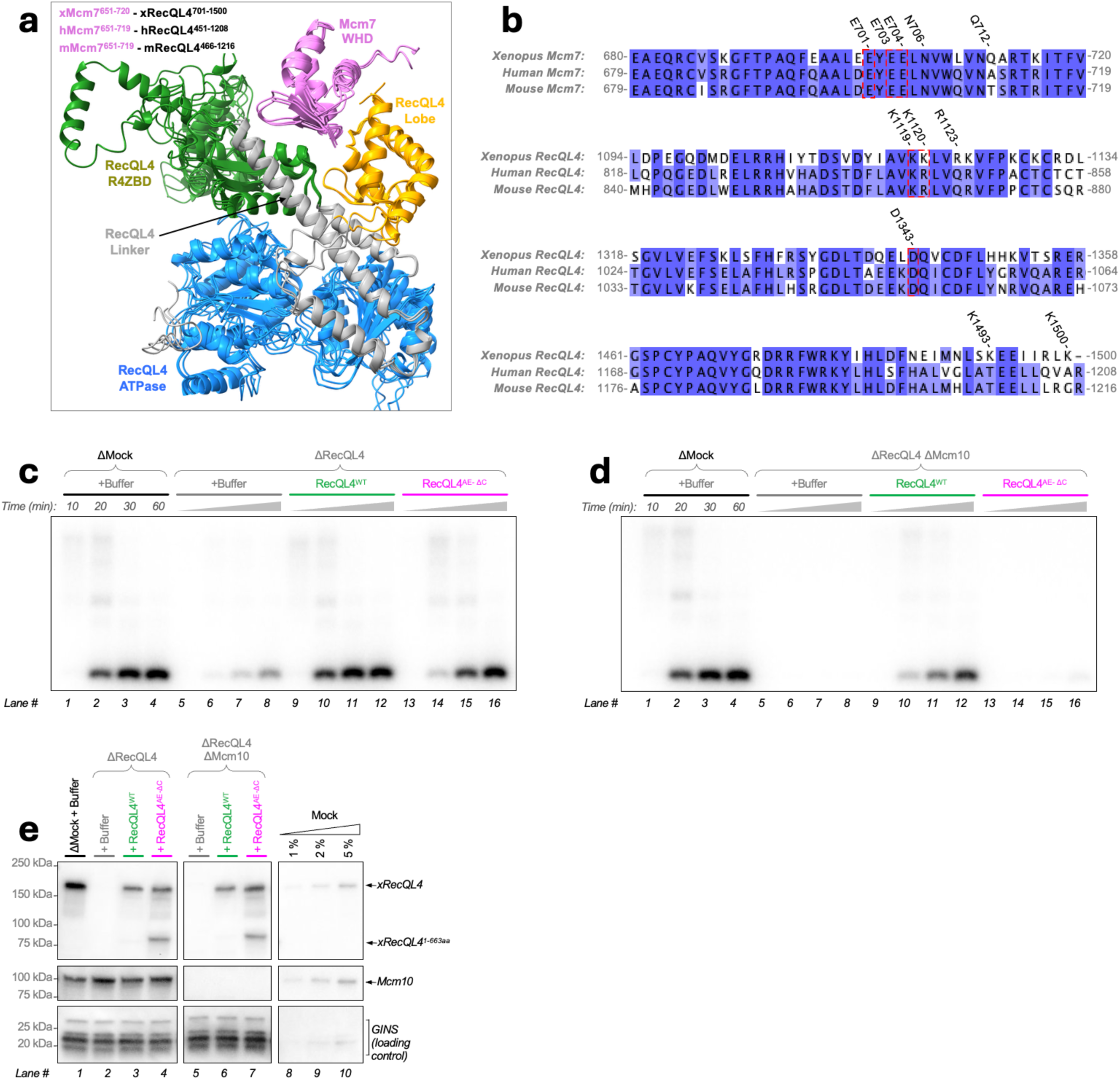
**(a)** AlphaFold3-predicted structures of Mcm7’s winged-helix-domain (WHD, magenta) bound to the structured portion of RecQL4 for frog (x), human (h), and mouse (m). The top models for each organism are shown overlayed. Different RecQL4 domains are colored to match the colors in the domain diagram from Fig. 6b. **(b)** Multiple-sequence alignment of frog/human/mouse Mcm7 and RecQL4 highlighting regions and residues involved in the AlphaFold3-predicted interaction (generated using Clustal Omega^5^). **(c)** Raw autoradiograph corresponding to data shown in Fig. 6d. **(d)** Raw autoradiograph corresponding to data shown in Fig. 6e. **(e)** Immunoblots of extract samples taken from reactions shown in panels **(c)** and **(d)**. GINS serves as a loading control.

**Extended Data Figure 7.**
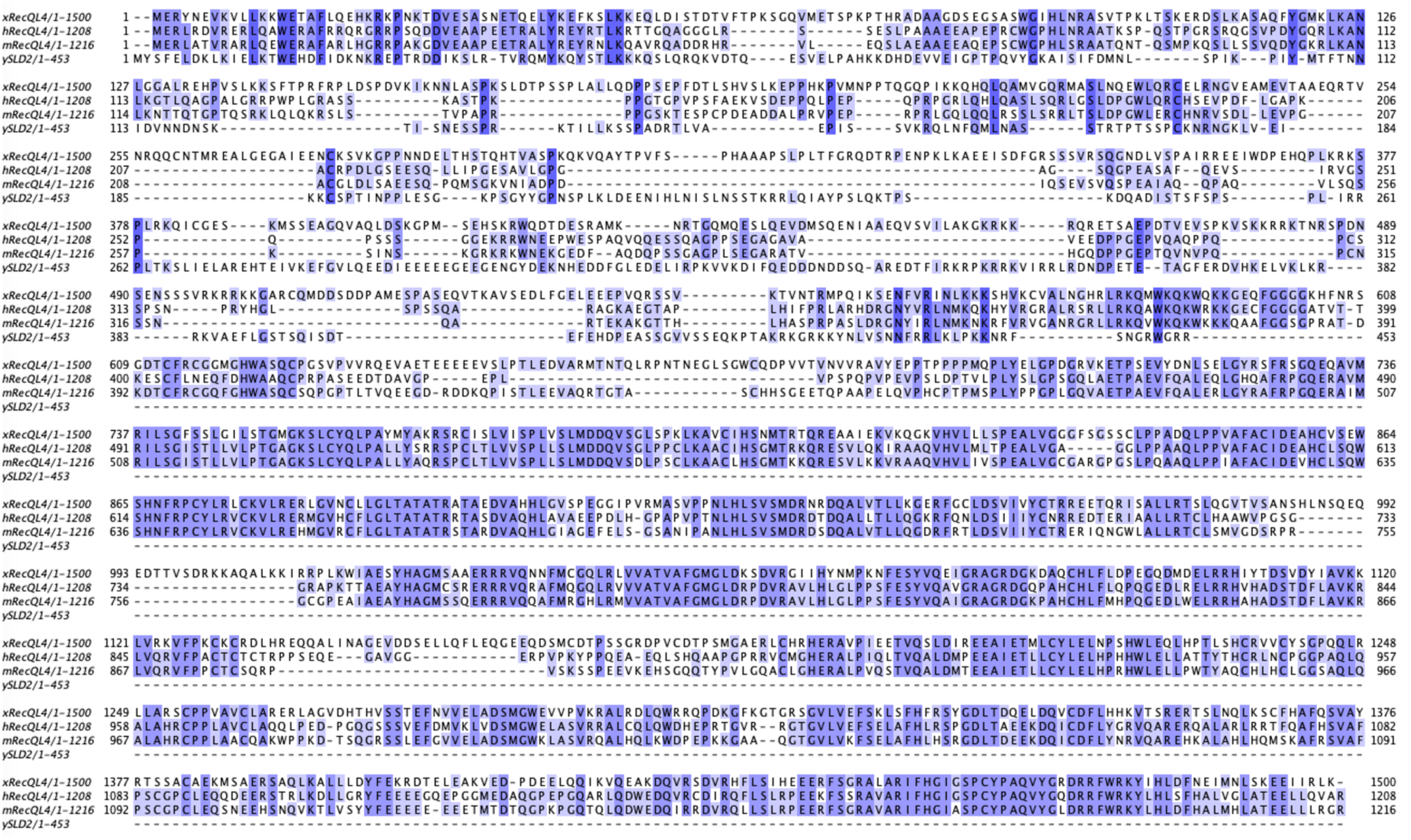
Full-length multiple-sequence alignment of *Saccharomyces cerevisiae* (y) Sld2 versus RecQL4 from *Xenopus laevis* (x), human (h), mouse (m) generated using Clustal Omega^5^. It highlights high sequence conservation among metazoa, and poor sequence conservation between metazoa and yeast.

## SUPPLEMENTARY INFORMATION

### Lead contact

Further information and requests for resources and reagents should be directed to and will be fulfilled by the lead contact, Gheorghe Chistol.

### Materials availability

All newly generated reagents generated in this study are available upon request from the lead contact.

### Data and code availability

- All data reported in this paper will be shared by the lead contact upon request.
- This paper does not report any original code.
- Any additional information required to reanalyze the data reported in this paper is available from the lead contact upon request.

### Xenopus Laevis Model System

Eggs from adult lab bred female *Xenopus laevis* frogs (Xenopus 1, Cat# 4800) were used to prepare egg extracts. Testes from adult lab bred male *Xenopus laevis* frogs (Xenopus 1, Cat# 4293) were used to prepare sperm chromatin. All animals were healthy, not subjected to previous procedures, and no animal husbandry was performed. Frogs were housed at the Boswell Small Amphibians Research Facility at Stanford School of Medicine in compliance with Institute Animal Care and Use Committee (IACUC) regulations and all experiments involving frogs were approved by the Stanford University APLAC (Administrative Panel on Laboratory Animal Care).

### Insect Cell Lines

Sf9 cells (Expression Systems, Cat# 94-001S) and Tni cells (Expression Systems, Cat# 94-002S) were cultured in ESF 921 insect cell culture medium (Expression Systems, Cat# 96-001-01) at 27°C for baculovirus production and protein overexpression.

### Bacterial Cell Lines

DH10EMBacY (Geneva Biotech, Cat# DH10EMBacY), DH5α (New England Biorabs (NEB), Cat# C2987H), and Rosetta (DE3) pLysS (Novagen, Cat# 70956-4) bacteria were cultured in LB broth (Thermo Fisher Scientific, Cat# BP9723) at 37ᵒC for plasmid production and protein overexpression.

### IVTT System

IVTT reactions were performed using TnT SP6 High-Yield Wheat Germ Protein Expression System (Promega, Cat# L3261) according to the manufacturer’s instructions. Proteins were not purified after their production by IVTT. The IVTT reaction was directly added replication reactions as the source of a given protein. The optimal concentration of the template plasmid was titrated for each protein.

### Preparation of DNA Substrates

All single molecule experiments used doubly tethered λ phage DNA. The 48.5 kb DNA substrate as purchased from NEB (#N3011S) has 12-nt 5’ ssDNA overhangs on both ends. We used the Klenow Fragment (3′ to 5′ exo-) polymerase (NEB, Cat# M0212S) to fill in the overhangs with biotin-dCTP (Thermo Fisher Scientific, Cat# 19518018) and biotin-dGTP (PerkinElmer, Cat# NEL541001EA). The product was separated via electrophoresis and electro-eluted from the gel before use in tethering. A step-by-step detailed protocol for preparing biotinylated λ DNA is available in our dedicated methods paper^1^.

### Xenopus Egg Extract Preparation

*Xenopus laevis* egg extracts were prepared following published protocols^2,3^. Frogs were primed with 75 IU of human chorionic gonadotropin (CHORULON) (Merck, Cat# 133754) 5-7 days before extract preparation. To induce ovulation, 660 IU of CHORULON was injected 20-21 hr before extract preparation. High-speed supernatant (HSS) was prepared from 5-6 adult female frogs, and nucleoplasmic extract (NPE) was prepared from 15-20 frogs. Sperm chromatin was purified from adult male frog testes and used for NPE preparation.

### Ensemble Replication Assay

To validate antibodies and recombinant proteins, ensemble DNA replication experiments were performed following a detailed protocol described in a recent methods article^1^. Since all proteins of interest (POIs) in this study are dispensable for Mcm2-7 double-hexamer loading, all licensing reactions were performed with HSS depleted of the POI.

### Protein Co-Immunoprecipitation Assay

For the co-immunoprecipitation experiment with recombinant FLAG-RecQL4 and untagged Mcm10, untagged *Xenopus laevis* Mcm10 was expressed in IVTT from [pRT81]pSP64-xMcm10 at 25°C for 2 hours. Anti-FLAG M2 Magnetic Beads (Millipore #M8823) were pelleted and washed three times with binding buffer (25 mM HEPES-KOH pH 7.5, 150 mM NaCl, 5% glycerol, 1 mM DTT) and aliquots with 3 μL packed beads were made. As a negative control reaction, 6 μL of 300 nM 3xFLAG peptide (APExBio, Cat# A6001) and 3 μL of IVTT reaction containing 600 nM Mcm10 were added to 3 μL of washed anti-FLAG M2 Magnetic Beads. To co-immunoprecipitate Mcm10 with FLAG-RecQL4, 6 μL of 300 nM FLAG-RecQL4 and 3 μL of IVTT reaction containing 600 nM Mcm10 were added to 3 μL of washed anti-FLAG M2 Magnetic Beads. These mixtures were incubated for 2 hours at 25°C with gentle rotation. Next, beads were washed three times with IP buffer (25 mM HEPES-KOH pH 7.5, 100 mM KCl, 10% glycerol, 0.1 mg/mL BSA, 1 mM DTT). Finally, beads were resuspended in 6 μL of IP buffer and transferred to a fresh tube. Bound proteins were eluted by boiling beads in 12 μL 1x Laemmli buffer (50mM Tris-HCl pH 6.8, 2% SDS, 10% glycerol, 0.1% bromophenol blue, 5% β-mercaptoethanol). To carefully quantify the co-immunoprecipitation efficiency, recombinant protein standards were loaded alongside eluted samples on the acrylamide gel.

For the co-immunoprecipitation experiment with recombinant FLAG-RecQL4 and untagged CMG complex, anti-FLAG M2 Magnetic Beads were pelleted and washed three times with high-salt binding buffer (25 mM HEPES-KOH pH 7.5, 500 mM NaCl, 5% glycerol, 1 mM DTT) and aliquots with 3 μL of packed beads were made. 3 μL of washed anti-FLAG M2 Magnetic Beads were incubated with 9 μL of 200 nM 3xFLAG peptide, 200 nM FLAG-RecQL4^WT^, or 200 nM FLAG-RecQL4^AE-ΔC^ for 2 hours at 4°C with gentle rotation. Beads were washed twice with high-salt binding buffer, and then, washed twice with IP buffer. 3 μL of pre-incubated packed beads were incubated with 6 μL of 100 nM CMG complex purified from Tni cells infected with [pGC299-b2] for 1 hour at 4°C with gentle rotation. 9 μL of IP reaction was overlaid to 400 μL of IP-0.5M Sucrose cushion (25 mM HEPES-KOH pH 7.5, 100 mM KCl, 0.5 M Sucrose, 0.1 mg/mL BSA, 1 mM DTT) in a 0.75-mL tube. Beads were immediately pelleted by centrifugation at 1,2000 xg for 2 min. Beads were resuspended in 6 μL of IP buffer and transferred to a fresh tube. Bound proteins were eluted by boiling beads in 12 μL 1x Laemmli buffer (50mM Tris-HCl pH 6.8, 2% SDS, 10% glycerol, 0.1% bromophenol blue, 5% β-mercaptoethanol). To carefully quantify the co-immunoprecipitation efficiency, recombinant protein standards were loaded alongside eluted samples on the acrylamide gel.

### Western Blotting and Coomasie Staining

All samples were boiled in 1x Laemmli buffer and run on 4-15% Mini-PROTEAN TGX Precast Protein Gels (Bio-Rad #4561086) using Tris-Glycine-SDS Running Buffer (Boston BioProducts #BP-150). EZ-Run Prestained Rec Protein Ladder (Fisher Scientific #BP36031) or Precision Plus Protein Dual Color Standard (Bio-Rad #1610374EDU) was run alongside samples to infer protein band sizes. For Coomasie staining, protein gels were incubated with Optiblot Blue InstantBlue Coomassie Protein Stain (Abcam #ab119211) for at least 1 hour at room temperature with gentle shaking. For immunoblotting, proteins were transferred from the gel onto a PVDF membrane (VWR #10061-492) in transfer buffer (Fisher Scientific #NC9297917). Membranes were blocked in 1x PBST (1x Phosphate Buffered Saline (PBS) (Fisher Scientific, Cat# NC9140736), 0.05% Tween 20) containing 5% (w/v) nonfat milk for 1 hour at room temperature with gentle shaking. Following a wash in 1xPBST, the membrane was incubated with primary antibodies diluted in 1x PBST containing 1% (w/v) BSA overnight at 4ᵒC with gentle shaking. Membranes were washed three times with 1x PBST and incubated with secondary antibody diluted in 1x PBST + 5% nonfat milk for 1 hour at room temperature with gentle shaking. Membranes were washed three times with 1x PBST, developed using SuperSignal West Pico PLUS Chemiluminescent Substrate (Fisher Scientific, Cat# PI34577), and imaged using Azure c600 (Azure Biosystems).

### Single Molecule Replication Assay

All single molecule experiments were conducted following a previously published protocol^1^, except for the following modifications. In the λ DNA tethering step, 20 pM of streptavidin^AF647^ (Thermo Fisher Scientific, Cat# S21374) was added to the streptavidin solution. Streptavidin^AF647^ spots were used as a fiducial marker for focusing immediately before starting imaging. After λ DNA tethering, DNA was licensed in HSS for 10 minutes. In the original KEHRMIT protocol, after a brief pulse of replication initiation at high GINS^AF647^ concentration (∼200 nM), the flow cell was flushed with GINS-depleted extract to prevent further origin firing and to minimize background^1,4^. In this study, Replication Mix (10 µL of NPE, 10 µL of HSS, 6 µL of ELB-Sucrose, 1 µL of 600 ng/µL pBlueScript carrier plasmid DNA, 1 µL of ATP regeneration Mix (67 mM ATP, 0.17 mg/mL creatine phosphokinase, 0.67 M phosphocreatine), 1 µL of 60 µM Fen1^mKikGR^, and 1 µL of 150-300 nM fluorescently labeled protein of interest) was flowed into the microfluidic chamber. The reaction was then imaged continuously (10 sec/frame) without washing out the free fluorescently labeled protein. Thus, fluorescent protein (5-10 nM) was present in the replication reaction during the entire duration of the experiment. Optimal replication efficiency was achieved when the single-molecule replication reaction contained equal volumes of NPE, HSS, and buffer^5^. The concentrations of proteins examined in this study were estimated using quantitative western blots (see table below).

| *Protein of Interest* | **Concentration in HSS (nM)** | **Concentration in NPE (nM)** | **Concentration in the Replication Reaction (nM)** |
| --- | --- | --- | --- |
| **RecQL4** | ~80 nM | ~700 nM | ~250 nM |
| **Mcm10** | ~20 nM | ~300 nM | ~100 nM |

For two-color imaging, Fen1^mKikGR^ (488 nm laser excitation) and the POI labeled with Alexa Fluor 647 (640 nm excitation) were used. The following settings were used for imaging Alexa Fluor 647: TIRF arm set to 9000 (∼67°), 500 ms exposure, 10 s/frame, 20% 640 nm laser power (∼250 µW power out of the objective, ∼2.5 W/cm^2^ power density), 100 EM Gain. Fen1^mKikGR^ was imaged using the following settings: TIRF arm set to 9000 (∼67°), 200 ms exposure, 10 s/frame, 8% 488 nm laser power (∼70 µW power out of the objective, ∼0.7 W/cm^2^ power density), 100 EM Gain. Given these exposure durations, 4 fields of view (FOVs) could be sequentially imaged during the 10 s/frame interval.

### Cloning and Plasmids

To prepare an antigen consisting of *Xenopus laevis* Mcm10 amino acids 128-240 with a His6 tag at the C-terminus, the geneblock GC601gb was inserted into pET-28b(+) linearized via digestion with NcoI and SacI via Gibson Assembly, yielding the plasmid [pLS002] pET-28b(+)-xMcm10^128-240^-His6.

A plasmid encoding the full-length frog *mcm10* gene was a generous gift from Justin Sparks. It was used as a PCR template to clone [pLS049]pFastBac-His6-FLAG-HRV3C-xMcm10.

To express Mcm10 in IVTT, the *mcm10* gene was PCR amplified from [pLS049] using oLAS006 and oLAS007. The resulting untagged *mcm10* gene product was then digested with SalI and SacI and cloned into the SalI-SacI region of pSP64 Poly(A), resulting in the plasmid [pLAS003]pSP64-xMcm10. To add a 10aa linker, a SNAP tag, and a FLAG tag to the C-terminus of Mcm10, the *mcm10* gene fragment containing a 10aa linker sequence was PCR amplified from [pLAS003] using primers oRT117 and oRT190, while the tandem 10aa-SNAP-FLAG linker sequence was PCR amplified from [pRT065]pSP64-xDONSON-SNAP-FLAG using primers oRT093 and oRT094. The two resulting PCR products were fused by overlap extension PCR using primers oRT117 and oRT094. The resulting xMcm10-10aa-SNAP-FLAG fragment was digested with SalI-HF and cloned into the SalI-SmaI site of pSP64 Poly(A), yielding the plasmid [pRT079]pSP64-xMcm10-SNAP-FLAG.

To express N-terminally FLAG tagged RecQL4, the untagged RecQL4 fragment was digested from [pRT025]pFastBac-xRecQL4 by NdeI and HindIII-HF, and inserted into the NdeI-HindIII site of [pRT027]pFastBac-FLAG-HRV3C-xRecQL4-10aa-sort-His6, yielding the plasmid [pRT074]pFastBac-FLAG-HRV3C-xRecQL4.

To express N-terminally tandem V5-SNAP-10aa tagged RecQL4 in IVTT, the NdeI-PvuI region of [pRT033]pSP64-V5-SNAP-xRecQL4-FLAG was substituted with the NdeI-PvuI region of [pRT030]pSP64-xRecQL4, yielding the plasmid [pRT082]pSP64-FLAG-SNAP-xRecQL4.

To clone the plasmid needed for expressing FLAG-HRV3C-RecQL4^1-663^, a fragment of the *recql4* gene was PCR amplified from pRT027 using phosphorylated primers oRT118 and oRT188. The resulting PCR product was circularized by ligation, yielding the plasmid [pRT077]pFastBac-FLAG-HRV3C-xRecQL4^1-663^.

To clone the plasmid needed for expressing FLAG-HRV3C-RecQL4^AE-ΔC^, the gene fragments were amplified by PCR using primer pairs, oRT117 and oRT195, and oRT196 and oRT197, using pRT033 as a template. The two PCR fragments were fused by overlap-extension PCR with primers oRT117 and oRT197. The FLAG-HRV3C-xRecQL4^AE-ΔC^ fragment was digested with ApaI and XbaI and cloned into the same sites in pRT074 resulting in [pRT093]pFastBac-FLAG-HRV3C-xRecQL4^AE-ΔC^.

To clone the plasmid needed for expressing FLAG-HRV3C-RecQL4^664-1500^, a fragment of the *recql4* gene was PCR amplified from pRT074 using phosphorylated primers oRT198 and oRT199. The resulting PCR product was circularized by ligation, yielding the plasmid [pRT094]pFastBac-FLAG-HRV3C-xRecQL4^664-1500^.

To clone the plasmid needed for expressing FLAG-HRV3C-RecQL4^664-1500(D856A^), a fragment of the ATPase-dead *recql4* gene was PCR amplified from [pRT076]pFastBac-FLAG-HRV3C-xRecQL4^D856A^ using phosphorylated primers oRT198 and oRT199. The resulting PCR product was circularized by ligation, yielding the plasmid [pRT095]pFastBac-FLAG-HRV3C-xRecQL4^664-1500(D856A)^.

To express V5-SNAP-RecQL4^1-663^ in IVTT, a truncated *recql4* gene fragment was PCR amplified from pRT033 using phosphorylated primers oRT117 and oRT118. The resulting PCR product was digested by BsrGI-HF and inserted into the BsrGI-SmaI site of pSP64 Poly(A), yielding the plasmid [pRT049]pSP64-V5-SNAP-xRecQL4^1-663^.

To express V5-SNAP-RecQL4^664-1500^ in IVTT, a truncated *recql4* gene fragment was PCR amplified from pRT025 using primers oRT86 and oRT200, while the V5-SNAP-10aa sequence was PCR amplified from the gene block gRT88 using primers oRT089 and oRT090. The two resulting PCR products were fused by overlap extension PCR using primers oRT089 and oRT086. The resulting V5-SNAP-xRecQL4^664-1500^ fragment was digested with BmgBI and XmaI and cloned into the BmgBI-XmaI site of pRT082, yielding the plasmid [pRT085]pSP64-V5-SNAP-xRecQL4^664-1500^.

To express frog CMG in insect cells from a single baculovirus, we used the MultiBac^6^ cloning strategy to generate [pGC299] – a single plasmid encoding all 11 subunits of Xenopus laevis CMG as previously described^7,8^ with one difference: TwinStrep-HRV3C-Mcm3 was cloned in [pGC299] instead of FLAG-Mcm3 in [pGC187].

### Antibody Preparation

Custom polyclonal antibodies were generated as previously described^1^. Briefly, the His6 tagged antigen sequence was cloned into pET28b(+) (see Cloning and Plasmids). The antigen was expressed in Rosetta (DE3) pLysS (Novagen, Cat# 70956-4) upon induction with 0.5-1.0 mM IPTG for 3 hrs at 37°C or overnight at 16°C. Soluble antigens were purified in native buffer, whereas insoluble antigens were purified in denaturing buffer (see table and ‘Recombinant Protein Expression and Purification’ below). In most cases, the Ni-NTA eluate (still containing 400 mM imidazole) was used to immunize rabbits. For some soluble antigens, additional chromatography was used to polish the protein preparation. All antibodies used in this study were raised by Cocalico Biologicals.

All antibodies used in this study were affinity purified. 1-5 mg of purified antigen was cross-linked to 1 mL of AminoLink Plus Coupling Resin (Fisher Scientific, Cat# PI20501) according to the manufacturer’s protocol. 10-30 mL of serum was incubated with the affinity resin for 30 minutes at room temperature. The resin was washed twice with 10 mL of 1x PBS + 500 mM NaCl, and twice again with 10 mL of 1x PBS. Next, antibodies bound to the affinity resin were eluted 5-6 times with 1.0 mL of 200 mM glycine-HCl pH2.5. Each elution was collected into an tube containing 0.1 mL of 1M Tris-HCl pH9.0, which immediately neutralized the glycine. Eluted fractions were pooled and dialyzed against 1 L of 1x TBS (50 mM Tris-HCl pH8.0, 150 mM NaCl) + 10% (w/v) sucrose. The IgG concentration was measured using a Nanodrop, normalized to 1 mg/mL via dilution or spin-concentration, aliquots were flash frozen in liquid nitrogen, and stored at -80°C.

| Target Protein<br>(Rabbit #) | Antigen | Plasmid<br>(expression) | Antigen<br>Purification | Depletion<br>Condition<br>µg IgG per<br>µL extract | Primary<br>Antibody<br>Concentration<br>for WB |
| --- | --- | --- | --- | --- | --- |
| Anti-xGINS<br>(SU289) | whole GINS complex<br>Psf2-10aa-sort-His6 | pGC127<br>(insect) | Native | 2:1 | 0.2 µg/mL |
| Anti-Cdc45<br>(SU294) | (1-361 aa)-His6 | pGC283<br>(bacteria) | Denaturing | 1:1 | 0.2 µg/mL |
| Anti-xRecQL4<br>(SU307) | (56-165 aa)-His6 | pSEB014<br>(bacteria) | Denaturing | 1:1 | 0.2 µg/mL |
| Anti-xRecQL4<br>(SU305) | (1145-1319 aa)-His6 | pSEB011<br>(bacteria) | Denaturing | N/A | 0.2 µg/mL |
| Anti-xDONSON<br>(SU351) | His6-(155-579 aa) | pRT057<br>(bacteria) | Native | 1:1 | 0.2 µg/mL |
| Anti-xMcm10<br>(SU255) | (128-240 aa)-His6 | pLS002<br>(bacteria) | Native | 1:1 | 0.2 µg/mL |
| Anti-xMcm7<br>(SU347) | His6-(400-720 aa) | pRT053<br>(bacteria) | Denaturing | N/A | 0.2 µg/mL |

### Recombinant Protein Expression, Purification

Previously published protocols were used to purify recombinant Cdc45^9^, DONSON^9^, Geminin^11^, GINS^4^, RecQL4^9^, and Fen1-mKikGR^13^.

A bacmid was generated by transforming DH10EMBacY cells with [pRT074]pFastBac1-FLAG-HRV3C-xRecQL4^WT^, [pRT077] pFastBac-FLAG-HRV3C-xRecQL4^1-663^, [pRT093] pFastBac-FLAG-HRV3C-xRecQL4^AE-ΔC^, [pRT094] pFastBac-FLAG-HRV3C-xRecQL4^664-1500^, [pRT095] pFastBac-FLAG-HRV3C-xRecQL4^664-1500(D856A)^ and the resulting bacmid was purified with the ZR BAC DNA Miniprep Kit (Zymo Research, Cat# D4049). The baculovirus was amplified in three stages (P1, P2, P3) in Sf9 cells.

RecQL4^1-663^, RecQL4^664-1500^, and RecQL4^664-1500(D856A)^ were purified following the same protocol^9^ as RecQL4^WT^ and RecQL4^D856A^. Briefly, proteins were expressed in 250 mL of Sf9 suspension culture by infection with the P3 baculovirus for 48 hrs. Cells were harvested, washed with PBS, pelleted by centrifugation, and flash frozen in liquid nitrogen. Cell pellets were thawed and resuspended in four volumes of buffer L (25 mM HEPES pH7.5, 300 mM NaCl, 10% glycerol, 1 mM DTT) containing 1X Pierce Protease Inhibitor EDTA-free (ThermoFisher Scientific, Cat# A32965). Lysate was cleared by ultracentrifugation at 35,000 rpm for 40 min at 4°C in a Beckman Type 70 Ti rotor (Beckman Coulter). Cleared lysate was incubated with 200 µL of ANTI-FLAG M2 Affinity Gel for 1 hr at 4°C. The protein was eluted from the resin by cutting the HRV3C site with 50 µg/mL HRV3C protease in buffer L for 2 hours at 4°C. The eluted RecQL4 was separated from the HRV3C protease via size exclusion chromatography on a Superdex 200 Increase 10/300 GL column (Cytiva, Cat# 28990944). Peak fractions were pooled, concentrated, and snap frozen in LN_2_ as small aliquots, and stored at -80°C.

FLAG-RecQL4^WT^ and FLAG-RecQL4^AE-ΔC^ were purified as follows: Proteins were expressed in 250 mL of Sf9 suspension culture by infection with the P3 baculovirus for 48 hrs. Cells were harvested, washed with PBS, pelleted by centrifugation, and flash frozen in liquid nitrogen. Cell pellets were thawed and resuspended in four volumes of buffer F (25 mM HEPES pH7.5, 500 mM NaCl, 10% glycerol, 1 mM DTT) containing 1X Pierce Protease Inhibitor EDTA-free. Lysate was cleared by ultracentrifugation at 35,000 rpm for 40 min at 4°C in a Beckman Type 70 Ti rotor. Cleared lysate was incubated with 200 µL of ANTI-FLAG M2 Affinity Gel for 1 hr at 4°C. The protein was eluted from the FLAG-M2 resin with 100 µg/mL 3X FLAG peptide (APExBio, Cat# A6001) in buffer F containing 1X Pierce Protease Inhibitor EDTA-free. The eluted protein was clarified via size exclusion chromatography on a Superdex 200 Increase 10/300 GL column. The peak fractions were pooled and concentrated, snap frozen in liquid nitrogen as small aliquots, and stored at -80°C.

Recombinant frog CMG complex was purified as follows: A bacmid was generated by transforming DH10EMBacY cells with pGC299 and the resulting bacmid was purified with the ZR BAC DNA Miniprep Kit. The baculovirus was amplified in three stages (P1, P2, P3) in Sf9 cells. The protein was expressed in 1 L of Tni suspension culture by infection with the P3 baculovirus for 48 hrs. Cells were harvested, washed with PBS, pelleted by centrifugation, and flash frozen in liquid nitrogen. Cell pellets were thawed and resuspended in four volumes of buffer S (25 mM HEPES pH7.5, 300 mM NaOAc, 5 mM MgCl_2_, 10% glycerol, 0.1% Tween-20, 1 mM DTT) containing 1X Pierce Protease Inhibitor EDTA-free. Lysate was cleared by ultracentrifugation at 35,000 rpm for 40 min at 4°C in a Beckman Type 70 Ti rotor. Cleared lysate was incubated with 350 μL of StrepTactin XT 4Flow resin (IBA Lifesciences, Cat#2-5010-002) for 1 hr at 4°C. The CMG complex was eluted from the resin by cutting the HRV3C site with 50 µg/mL HRV3C protease in buffer S for 2 hours at 4°C. The eluate was dilute threefold in dilution buffer (25 mM HEPES pH7.5, 3 mM MnCl_2_, 10% glycerol, 1 mM DTT), and loaded onto 1 mL of Nuvia S resin (Biorad, Cat#1560311) packed into a 10-mL Poly-Prep chromatography column (Biorad, Cat#7311550) (final salt concentration was 100 mM NaOAc). The flowthrough from the Nuvia S column was incubated with 2.5 units/µL of Lambda protein phosphatase (NEB Cat#P0753S) for 30 min at room temperature, and loaded onto a Capto HiRes Q 5/50 (Cytive, Cat#29275878). The CMG complexes bound to the Capto HiRes Q column were eluted over a gradient of 100-700 mM KCl in buffer A (25 mM HEPES pH7.5, 1 mM EDTA, 1 mM EGTA, 10% glycerol, 0.02% Tween-20, 1 mM DTT). Fractions corresponding to the CMG peak were collected and diluted threefold in buffer A (final salt concentration was 150 mM KCl). The diluted CMG pool was concentrated by binding to 30 µL of Nuvia Q resin (Biorad, Cat#1560411) for 30 min at 4°C and then eluting with 70 µL of 600 mM KCl in buffer A. The eluates were flash-frozen in 2.5-µL aliquots in liquid nitrogen, and stored at -80°C.

The native purification of antigens was performed as follows. The Rosetta (DE3) pLysS cell pellet overexpressing the His6-tagged antigen was thawed in Lysis buffer containing 1 mM PMSF, and sonicated. The lysate was cleared by ultracentrifugation at 35,000 rpm for 40 min at 4°C in a Beckman Type 70 Ti rotor. Cleared lysate was incubated with 1 mL of Ni-NTA resin for 1 hr at 4°C. The protein was eluted with 400 mM imidazole in Lysis buffer containing 1 mM PMSF.

The denaturing purification of antigens was performed as follows. The Rosetta (DE3) pLysS cell pellet overexpressing His6-tagged antigen was thawed in Denaturing Lysis buffer (50 mM HEPES pH7.5, 500 mM NaCl, 10 mM imidazole, 8 M Urea) containing 1 mM PMSF, and sonicated. The lysate was cleared by ultracentrifugation at 35,000 rpm for 40 min at room temperature in a Beckman Type 70 Ti rotor. Cleared lysate was incubated with 1 mL of Ni-NTA resin for 1 hr at room temperature. The protein was eluted with 400 mM imidazole in denaturing elution buffer (50 mM HEPES pH7.5, 500 mM NaCl, 6 M Urea, 1 mM PMSF).

### Quantification of Proteins Expressed in IVTT

The concentration of proteins expressed in IVTT reactions was determined by semi-quantitative western blotting. Typically, 0.05 µL of the IVTT reaction expressing the protein of interest (POI) was loaded on an SDS-PAGE gel, and a dilution series (5-200 fmol) of purified recombinant POI was loaded alongside. After transfer onto a PVDF membrane, the membrane was blotted with specific antibody against the POI. The concentration of POI in the IVTT reaction was estimated by comparing its band intensity to those from the dilution series of purified proteins.

### Fluorescent Labeling using SNAP

After expressing RecQL4 or Mcm10 in the IVTT lysate, the reaction was kept on ice, and the concentration of the target protein was determined via quantitative western blotting. The SNAP labeling reagent was used in a two-fold molar excess relative to the target protein. SNAP-Surface Alexa Fluor 647 was added to the IVTT reaction containing the target protein and reacted overnight at 4°C with gentle end-over-end rotation. The final reaction was aliquoted, snap frozen in liquid nitrogen, and stored at -80°C. Fluorescent labeling efficiency was estimated by loading the IVTT reaction on a PAGE-Gel alongside a titration series of purified fluorescent protein standards (GINS-AF647). The labeled proteins were not purified after the SNAP-labeling reaction. The IVTT reaction was directly added replication reactions as the source of a given protein.

### Data Reporting

Details of the statistical analyses performed are indicated in the figure legends. Since some quantities measured in this study did not have symmetric distributions, we reported the median of the distribution. Unless otherwise specified in the figure captions, error bars represent the 95% confidence interval for the median. This was calculated using MATLAB’s built in bootstrapping function. The exact meaning of the number ‘N’ reported in each figure is indicated in the figure captions, it usually refers to the number of DNA molecules, the number of replication origins, or the number of binding events. We typically used N∼100 events to report the duration of an intrinsically stochastic event (for example the duration of a productive binding event). p-values were computed using the two-sample Kolmogorov-Smirnov test function built into MATLAB; ‘n.s.’ denotes not significant, ∗ = p value <0.05, ∗∗ = p value <0.01, ∗∗∗ = p value <0.001, ∗∗∗∗ = p value ≤0.0001.

### Reproducibility and Blinded Analysis

All experiments were repeated at least twice, with independent extract preparations. The data presented in the main figures comes from one of these preparations. Treatment series (for example Fig. 1g, Fig. 2b-e, Fig. 3b-e) were performed with the same extract for consistency. All the single-molecule data was analyzed in a blinded manner. Specifically, raw data from different experiments in the same treatment series (mock-depletion, Mcm10/RecQL4-depletion, recombinant protein add-back) were loaded into our custom-written data analysis package, the order of individual DNA molecules was scrambled, and the exact experiment corresponding to each DNA molecule was unknown to the user performing the analysis. The results were then unscrambled using custom MATLAB code written in-house.

### Single-Molecule Data Analysis

Each replication initiation experiment imaged 4-6 fields of view (FOVs) over the course of 45 min with a resolution of 10 seconds/frame. Each FOV typically contained 50-200 DNA molecules stretched to 80-90% of their contour length. A suite of MATLAB scripts written in house were used to select and analyze data as described in our recent methods article^10^. Below we summarize the data analysis pipeline specific for replication initiation experiments.

### Selecting DNA Molecules

Regions of interest (ROIs) corresponding to individual DNA molecules were selected manually based on the previews of DNA stained with SYTOX Green (these were acquired before the start of the experiment). Only well-separated DNA molecules were used in the analysis.

### Quantifying the Fen1 Signal

Custom MATLAB scripts written in house were used to generate kymograms corresponding to each selected DNA molecule. Origin firing was monitored by visualizing the binding of Fen1^mKikGR^ to nascent DNA. To measure fork speed, the raw Fen1 signal was automatically thresholded using OTSU’s method via a built-in MATLAB function. The OTSU threshold binary image was used to fit straight lines to the edges of the replication bubble, and the fork speed value was extracted from the slope of the line (accounting for pixel size, the time resolution of the experiment, and the extent to which DNA was stretched). To accurately determine the time when an origin fired, the raw Fen1 signal was integrated over time, and the integrated signal was fit by two straight line segments: the first segment represents the background noise before origin firing, while the second segment corresponds to a linear increase in Fen1 signal after the start of DNA synthesis. The intersection of the two segments corresponds to the time of origin firing. This estimate agrees well with the origin firing time prediction from the OTSU threshold analysis.

### Analyzing Mcm10^AF647^ and RecQL4^AF647^ Binding/Dissociation

To analyze the binding and dissociation of Mcm10/RecQl4, we employed the Generalized Likelihood Ratio Test (GLRT). GLRT was used to convert noisy single-molecule images into binary images that were then segmented into binding events. The GLRT algorithm uses a sliding window (we used a window with a 5-pixel radius) to determine whether the signal at a given pixel is consistent with noise or is significantly different from noise. This approach is superior to simple image thresholding because it accounts for the intrinsic properties of background noise. We adapted GLRT code written in MATLAB from a previous publication^12^. When analyzing productive/non-productive binding events, we did not score any events that lasted less than two frames because we could not be confident that they represented bona-fide binding.

### Global Protein Binding Analysis

To quantify the binding of RecQL4^AF647^ and Mcm10^AF647^ to the origins in mock versus depleted extracts, we devised a global binding analysis. First, ROIs with and without DNA were selected based on the SYTOX Green images. Next, kymograms were generated for both regions and the GLRT algorithm was employed to identify binding events (including single-frame events). The total duration of all binding events was added and normalized to DNA length (sec/kb) as shown in Extended Data Fig. 1f. Matched selections without DNA were used to subtract non-specific background binding (which may vary between experiments).

## REFERENCES

1. Di Micco, R., Fumagalli, M., Cicalese, A., Piccinin, S., Gasparini, P., Luise, C., Schurra, C., Garré, M., Giovanni Nuciforo, P., Bensimon, A., et al. (2006). Oncogene-induced senescence is a DNA damage response triggered by DNA hyper-replication. Nature 444, 638–642. 10.1038/nature05327.

2. Macheret, M., and Halazonetis, T.D. (2018). Intragenic origins due to short G1 phases underlie oncogene-induced DNA replication stress. Nature 555, 112–116. 10.1038/nature25507.

3. Dominguez-Sola, D., Ying, C.Y., Grandori, C., Ruggiero, L., Chen, B., Li, M., Galloway, D.A., Gu, W., Gautier, J., and Dalla-Favera, R. (2007). Non-transcriptional control of DNA replication by c-Myc. Nature 448, 445–451. 10.1038/nature05953.

4. Murat, P., Guilbaud, G., and Sale, J.E. (2024). DNA replication initiation drives focal mutagenesis and rearrangements in human cancers. Nat. Commun. 10.1038/s41467-024-55148-3.

5. Remus, D., Beuron, F., Tolun, G., Griffith, J.D., Morris, E.P., and Diffley, J.F.X. (2009). Concerted Loading of Mcm2-7 Double Hexamers around DNA during DNA Replication Origin Licensing. Cell 139, 719–730. 10.1016/j.cell.2009.10.015.

6. Evrin, C., Clarke, P., Zech, J., Lurz, R., Sun, J., Uhle, S., Li, H., Stillman, B., and Speck, C. (2009). A double-hexameric MCM2-7 complex is loaded onto origin DNA during licensing of eukaryotic DNA replication. Proc. Natl. Acad. Sci. U. S. A. 106, 20240–20245. 10.1073/pnas.0911500106.

7. Costa, A., and Diffley, J.F.X. (2022). The Initiation of Eukaryotic DNA Replication. Annu. Rev. Biochem. 91, 107–131. 10.1146/annurev-biochem-072321-110228.

8. Ilves, I., Petojevic, T., Pesavento, J.J., and Botchan, M.R. (2010). Activation of the MCM2-7 Helicase by Association with Cdc45 and GINS Proteins. Mol. Cell 37, 247–258. 10.1016/j.molcel.2009.12.030.

9. Kumagai, A., Shevchenko, A., Shevchenko, A., and Dunphy, W.G. (2010). Treslin Collaborates with TopBP1 in Triggering the Initiation of DNA Replication. Cell 140, 349– 359. 10.1016/j.cell.2009.12.049.

10. Hashimoto, Y., Sadano, K., Miyata, N., Ito, H., and Tanaka, H. (2023). Novel role of DONSON in CMG helicase assembly during vertebrate DNA replication initiation. EMBO J., 1–17. 10.15252/embj.2023114131.

11. Kumagai, A., and Dunphy, W.G. (2017). MTBP, the partner of Treslin, contains a novel DNA-binding domain that is essential for proper initiation of DNA replication. Mol. Biol. Cell 28, 2998–3012. 10.1091/mbc.E17-07-0448.

12. Kumagai, A., and Dunphy, W.G. (2020). Binding of the Treslin-MTBP Complex to Specific Regions of the Human Genome Promotes the Initiation of DNA Replication. Cell Rep. 32, 108178. 10.1016/j.celrep.2020.108178.

13. Xia, Y., Sonneville, R., Jenkyn-Bedford, M., Ji, L., Alabert, C., Hong, Y., Yeeles, J.T.P., and Labib, K.P.M. (2023). DNSN-1 recruits GINS for CMG helicase assembly during DNA replication initiation in Caenorhabditis elegans. Science (80-.). 4, 88–100. 10.1126/science.adi4932.

14. Lim, Y., Tamayo-Orrego, L., Schmid, E., Tarnauskaite, Z., Kochenova, O. V, Gruar, R., Muramatsu, S., Lynch, L., Schlie, A.V., Carroll, P.L., et al. (2023). In silico protein interaction screening uncovers DONSON’s role in replication initiation. Science (80-.). 3448. 10.1126/science.adi3448.

15. Kingsley, G., Skagia, A., Passaretti, P., Fernandez-Cuesta, C., Reynolds-Winczura, A., Koscielniak, K., and Gambus, A. (2023). DONSON facilitates Cdc45 and GINS chromatin association and is essential for DNA replication initiation. Nucleic Acids Res., 1–16. 10.1093/nar/gkad694.

16. Cvetkovic, M.A., Passaretti, P., Butryn, A., Reynolds-Winczura, A., Kingsley, G., Skagia, A., Fernandez-Cuesta, C., Poovathumkadavil, D., George, R., Chauhan, A.S., et al. (2023). The structural mechanism of dimeric DONSON in replicative helicase activation. Mol. Cell 83, 4017–4031.e9. 10.1016/j.molcel.2023.09.029.

17. Terui, R., Berger, S.E., Sambel, L.A., Song, D., and Chistol, G. (2024). Single-molecule imaging reveals the mechanism of bidirectional replication initiation in metazoa. Cell 187, 3992–4009.e25. 10.1016/j.cell.2024.05.024.

18. Douglas, M.E., Ali, F.A., Costa, A., and Diffley, J.F.X. (2018). The mechanism of eukaryotic CMG helicase activation. Nature 555, 265–268. 10.1038/nature25787.

19. Henrikus, S.S., Gross, M.H., Willhoft, O., Pühringer, T., Lewis, J.S., McClure, A.W., Greiwe, J.F., Palm, G., Nans, A., Diffley, J.F.X., et al. (2024). Unwinding of a eukaryotic origin of replication visualized by cryo-EM. Nat. Struct. Mol. Biol. 31. 10.1038/s41594-024-01280-z.

20. Langston, L.D., Mayle, R., Schauer, G.D., Yurieva, O., Zhang, D., Yao, N.Y., Georgescu, R.E., and O’Donnell, M.E. (2017). Mcm10 promotes rapid isomerization of CMG-DNA for replisome bypass of lagging strand DNA blocks. Elife 6, 1–21. 10.7554/eLife.29118.

21. Langston, L.D., and O’Donnell, M.E. (2019). An explanation for origin unwinding in eukaryotes. Elife 8, 1–17. 10.7554/eLife.46515.

22. Langston, L.D., Georgescu, R.E., and O’Donnell, M.E. (2023). Mechanism of eukaryotic origin unwinding is a dual helicase DNA shearing process. Proc. Natl. Acad. Sci. 120, 2017. 10.1073/pnas.2316466120.

23. Watase, G., Takisawa, H., and Kanemaki, M.T. (2012). Mcm10 plays a role in functioning of the eukaryotic replicative DNA helicase, Cdc45-Mcm-GINS. Curr. Biol. 22, 343–349. 10.1016/j.cub.2012.01.023.

24. Yeeles, J.T.P., Deegan, T.D., Janska, A., Early, A., and Diffley, J.F.X. (2015). Regulated eukaryotic DNA replication origin firing with purified proteins. Nature 519, 431–435. 10.1038/nature14285.

25. Van Deursen, F., Sengupta, S., De Piccoli, G., Sanchez-Diaz, A., and Labib, K. (2012). Mcm 10 associates with the loaded DNA helicase at replication origins and defines a novel step in its activation. EMBO J. 31, 2195–2206. 10.1038/emboj.2012.69.

26. Luong, T.T., and Bernstein, K.A. (2021). Role and regulation of the recql4 family during genomic integrity maintenance. Genes (Basel). 12. 10.3390/genes12121919.

27. Ahmed, S.M.Q., Sasikumar, J., Laha, S., and Das, S.P. (2024). Multifaceted role of the DNA replication protein MCM10 in maintaining genome stability and its implication in human diseases. Cancer Metastasis Rev. 43, 1353–1371. 10.1007/s10555-024-10209-3.

28. Baxley, R.M., and Bielinsky, A.K. (2017). Mcm10: A dynamic scaffold at eukaryotic replication forks. Genes (Basel). 8, 13–16. 10.3390/genes8020073.

29. Mo, D., Zhao, Y., and Balajee, A.S. (2018). Human RecQL4 helicase plays multifaceted roles in the genomic stability of normal and cancer cells. Cancer Lett. 413, 1–10. 10.1016/j.canlet.2017.10.021.

30. Balajee, A.S. (2021). Human RecQL4 as a Novel Molecular Target for Cancer Therapy. Cytogenet. Genome Res., 1–23. 10.1159/000516568.

31. Wohlschlegel, J.A., Dhar, S.K., Prokhorova, T.A., Dutta, A., and Walter, J.C. (2002). Xenopus Mcm10 binds to origins of DNA replication after Mcm2-7 and stimulates origin binding of Cdc45. Mol. Cell 9, 233–240. 10.1016/S1097-2765(02)00456-2.

32. Chadha, G.S., Gambus, A., Gillespie, P.J., and Blow, J.J. (2016). Xenopus Mcm10 is a CDK-substrate required for replication fork stability. Cell Cycle 15, 2183–2195. 10.1080/15384101.2016.1199305.

33. Im, J.S., Ki, S.H., Farina, A., Jung, D.S., Hurwitz, J., and Lee, J.K. (2009). Assembly of the Cdc45-Mcm2-7-GINS complex in human cells requires the Ctf4/And-1, RecQL4, and Mcm10 proteins. Proc. Natl. Acad. Sci. U. S. A. 106, 15628–15632. 10.1073/pnas.0908039106.

34. Kanke, M., Kodama, Y., Takahashi, T.S., Nakagawa, T., and Masukata, H. (2012). Mcm10 plays an essential role in origin DNA unwinding after loading of the CMG components. EMBO J. 31, 2182–2194. 10.1038/emboj.2012.68.

35. Kliszczak, M., Sedlackova, H., Pitchai, G.P., Streicher, W.W., Krejci, L., and Hickson, I.D. (2015). Interaction of RECQ4 and MCM10 is important for efficient DNA replication origin firing in human cells. Oncotarget 6, 40464–40479. 10.18632/oncotarget.6342.

36. Xu, X., Rochette, P.J., Feyissa, E.A., Su, T. V., and Liu, Y. (2009). MCM10 mediates RECQ4 association with MCM2-7 helicase complex during DNA replication. EMBO J. 28, 3005–3014. 10.1038/emboj.2009.235.

37. Im, J.S., Park, S.Y., Cho, W.H., Bae, S.H., Hurwitz, J., and Lee, J.K. (2015). RecQL4 is required for the association of Mcm10 and Ctf4 with replication origins in human cells. Cell Cycle 14, 1001–1009. 10.1080/15384101.2015.1007001.

38. Wu, Z., Wang, Y., Li, J., Wang, H., Tuo, X., and Zheng, J. (2022). MCM10 is a Prognostic Biomarker and Correlated With Immune Checkpoints in Ovarian Cancer. Front. Genet. 13, 1–12. 10.3389/fgene.2022.864578.

39. Mughal, M.J., Chan, K.I., Mahadevappa, R., Wong, S.W., Wai, K.C., and Kwok, H.F. (2022). Over-Activation of Minichromosome Maintenance Protein 10 Promotes Genomic Instability in Early Stages of Breast Cancer. Int. J. Biol. Sci. 18, 3827–3844. 10.7150/ijbs.69344.

40. Xu, X., Chang, C.W., Li, M., Liu, C., and Liu, Y. (2021). Molecular Mechanisms of the RECQ4 Pathogenic Mutations. Front. Mol. Biosci. 8, 1–11. 10.3389/fmolb.2021.791194.

41. Arora, A., Agarwal, D., Abdel-Fatah, T.M.A., Lu, H., Croteau, D.L., Moseley, P., Aleskandarany, M.A., Green, A.R., Ball, G., Rakha, E.A., et al. (2016). RECQL4 helicase has oncogenic potential in sporadic breast cancers. J. Pathol. 238, 495–501. 10.1002/path.4681.

42. Lyu, G., Su, P., Hao, X., Chen, S., Ren, S., Zhao, Z., Gong, Y., Liu, Q., and Shao, C. (2021). RECQL4 regulates DNA damage response and redox homeostasis in esophageal cancer. Cancer Biol. Med. 18, 120–138. 10.20892/j.issn.2095-3941.2020.0105.

43. Baxley, R.M., Leung, W., Schmit, M.M., Matson, J.P., Yin, L., Oram, M.K., Wang, L., Taylor, J., Hedberg, J., Rogers, C.B., et al. (2021). Bi-allelic MCM10 variants associated with immune dysfunction and cardiomyopathy cause telomere shortening. Nat. Commun. 12, 1–19. 10.1038/s41467-021-21878-x.

44. Schmit, M.M., Baxley, R.M., Wang, L., Hinderlie, P., Kaufman, M., Simon, E., Raju, A., Miller, J.S., and Bielinsky, A.K. (2024). A critical threshold of MCM10 is required to maintain genome stability during differentiation of induced pluripotent stem cells into natural killer cells. Open Biol. 14. 10.1098/rsob.230407.

45. Mace, E.M., Paust, S., Conte, M.I., Baxley, R.M., Schmit, M.M., Patil, S.L., Guilz, N.C., Mukherjee, M., Pezzi, A.E., Chmielowiec, J., et al. (2020). Human NK cell deficiency as a result of biallelic mutations in MCM10. J. Clin. Invest. 130, 5272–5286. 10.1172/JCI134966.

46. Siitonen, H.A., Kopra, O., Kääriäinen, H., Haravuori, H., Winter, R.M., Säämänen, A.M., Peltonen, L., and Kestilä, M. (2003). Molecular defect of RAPADILINO syndrome expands the phenotype spectrum of RECQL diseases. Hum. Mol. Genet. 12, 2837–2844. 10.1093/hmg/ddg306.

47. Siitonen, A.H., Sotkasiira, J., Biervliet, M., Benmansour, A., Capri, Y., Cormier-Daire, V., Crandall, B., Hannula-Jouppi, K., Hennekam, R., Herzog, D., et al. (2009). The mutation spectrum in RECQL4 diseases. Eur. J. Hum. Genet. 17, 151–158. 10.1038/ejhg.2008.154.

48. Walter, J., Sun, L., and Newport, J. (1998). Regulated chromosomal DNA replication in the absence of a nucleus. Mol. Cell 1, 519–529. 10.1016/s1097-2765(00)80052-0.

49. Hoogenboom, W.S., Klein Douwel, D., and Knipscheer, P. (2017). Xenopus egg extract: A powerful tool to study genome maintenance mechanisms. Dev. Biol. 428, 300–309. 10.1016/j.ydbio.2017.03.033.

50. Berger, S., and Chistol, G. (2024). Visualizing the dynamics of DNA replication and repair at the single-molecule level. In Methods in Cell Biology (Elsevier Inc.), pp. 109–165. 10.1016/bs.mcb.2023.07.001.

51. Sparks, J.L., Chistol, G., Gao, A.O., Räschle, M., Larsen, N.B., Mann, M., Duxin, J.P., and Walter, J.C. (2019). The CMG Helicase Bypasses DNA-Protein Cross-Links to Facilitate Their Repair. Cell 176, 167–181.e21. 10.1016/j.cell.2018.10.053.

52. Loveland, A.B., Habuchi, S., Walter, J.C., and Van Oijen, A.M. (2012). A general approach to break the concentration barrier in single-molecule imaging. Nat. Methods 9, 987–992. 10.1038/nmeth.2174.

53. Sangrithi, M.N., Bernal, J.A., Madine, M., Philpott, A., Lee, J., Dunphy, W.G., and Venkitaraman, A.R. (2005). Initiation of DNA replication requires the RECQL4 protein mutated in Rothmund-Thomson syndrome. Cell 121, 887–898. 10.1016/j.cell.2005.05.015.

54. Matsuno, K., Kumano, M., Kubota, Y., Hashimoto, Y., and Takisawa, H. (2006). The N-Terminal Noncatalytic Region of Xenopus RecQ4 Is Required for Chromatin Binding of DNA Polymerase α in the Initiation of DNA Replication. Mol. Cell. Biol. 26, 4843–4852. 10.1128/mcb.02267-05.

55. Kaiser, S., Sauer, F., and Kisker, C. (2017). The structural and functional characterization of human RecQ4 reveals insights into its helicase mechanism. Nat. Commun. 8, 1–12. 10.1038/ncomms15907.

56. Warren, E.M., Vaithiyalingam, S., Haworth, J., Greer, B., Bielinsky, A.K., Chazin, W.J., and Eichman, B.F. (2008). Structural Basis for DNA Binding by Replication Initiator Mcm10. Structure 16, 1892–1901. 10.1016/j.str.2008.10.005.

57. Mayle, R., Langston, L., Molloy, K.R., Zhang, D., Chait, B.T., and O’Donnell, M.E. (2019). Mcm10 has potent strand-annealing activity and limits translocase-mediated fork regression. Proc. Natl. Acad. Sci. U. S. A. 116, 798–803. 10.1073/pnas.1819107116.

58. Xu, X., and Liu, Y. (2009). Dual DNA unwinding activities of the Rothmund-Thomson syndrome protein, RECQ4. EMBO J. 28, 568–577. 10.1038/emboj.2009.13.

59. Lõoke, M., Maloney, M.F., and Bell, S.P. (2017). Mcm10 regulates DNA replication elongation by stimulating the CMG replicative helicase. Genes Dev. 31, 291–305. 10.1101/gad.291336.116.

60. Padayachy, L., Ntallis, S.G., and Halazonetis, T.D. (2024). RECQL4 is not critical for firing of human DNA replication origins. Sci. Rep. 14, 1–12. 10.1038/s41598-024-58404-0.

61. Shin, G., Jeong, D., Kim, H., Im, J.S., and Lee, J.K. (2019). RecQL4 tethering on the pre-replicative complex induces unscheduled origin activation and replication stress in human cells. J. Biol. Chem. 294, 16255–16265. 10.1074/jbc.RA119.009996.

62. Thu, Y.M., and Bielinsky, A.K. (2013). Enigmatic roles of Mcm10 in DNA replication. Trends Biochem. Sci. 38, 184–194. 10.1016/j.tibs.2012.12.003.

63. Campos, L. V., Van Ravenstein, S.X., Vontalge, E.J., Greer, B.H., Heintzman, D.R., Kavlashvili, T., McDonald, W.H., Rose, K.L., Eichman, B.F., and Dewar, J.M. (2023). RTEL1 and MCM10 overcome topological stress during vertebrate replication termination. Cell Rep. 42, 112109. 10.1016/j.celrep.2023.112109.

64. Brosh, R.M., and Trakselis, M.A. (2019). Fine-tuning of the replisome: Mcm10 regulates fork progression and regression. Cell Cycle 18, 1047–1055. 10.1080/15384101.2019.1609833.

65. Alver, R.C., Zhang, T., Josephrajan, A., Fultz, B.L., Hendrix, C.J., Das-Bradoo, S., and Bielinsky, A.K. (2014). The N-terminus of Mcm10 is important for interaction with the 9-1-1 clamp and in resistance to DNA damage. Nucleic Acids Res. 42, 8389–8404. 10.1093/nar/gku479.

66. Wasserman, M.R., Schauer, G.D., O’Donnell, M.E., and Liu, S. (2019). Replication Fork Activation Is Enabled by a Single-Stranded DNA Gate in CMG Helicase. Cell 178, 600–611.e16. 10.1016/j.cell.2019.06.032.

67. Jones, M.L., Baris, Y., Taylor, M.R.G., and Yeeles, J.T.P. (2021). Structure of a human replisome shows the organisation and interactions of a DNA replication machine. EMBO J. 40, 1–23. 10.15252/embj.2021108819.

68. Martins, D.J., Di Lazzaro Filho, R., Bertola, D.R., and Hoch, N.C. (2023). Rothmund-Thomson syndrome, a disorder far from solved. Front. Aging 4, 1–13. 10.3389/fragi.2023.1296409.

69. Lindeboom, R.G.H., Vermeulen, M., Lehner, B., and Supek, F. (2019). The impact of nonsense-mediated mRNA decay on genetic disease, gene editing and cancer immunotherapy. Nat. Genet. 51, 1645–1651. 10.1038/s41588-019-0517-5.

70. Wang, L.L., Gannavarapu, A., Kozinetz, C.A., Levy, M.L., Lewis, R.A., Chintagumpala, M.M., Ruiz-Maldanado, R., Contreras-Ruiz, J., Cunniff, C., Erickson, R.P., et al. (2003). Association between osteosarcoma and deleterious mutations in the RECQL4 gene Rothmund-Thomson syndrome. J. Natl. Cancer Inst. 95, 669–674. 10.1093/jnci/95.9.669.

71. Cho, N.H., Cheveralls, K.C., Brunner, A.D., Kim, K., Michaelis, A.C., Raghavan, P., Kobayashi, H., Savy, L., Li, J.Y., Canaj, H., et al. (2022). OpenCell: Endogenous tagging for the cartography of human cellular organization. Science (80-.). 375. 10.1126/science.abi6983.

## EXTENDED DATA REFERENCES

1. Kaiser, S., Sauer, F., and Kisker, C. (2017). The structural and functional characterization of human RecQ4 reveals insights into its helicase mechanism. Nat. Commun. 8, 1–12. 10.1038/ncomms15907.

2. Terui, R., Berger, S.E., Sambel, L.A., Song, D., and Chistol, G. (2024). Single-molecule imaging reveals the mechanism of bidirectional replication initiation in metazoa. Cell 187, 3992–4009.e25. 10.1016/j.cell.2024.05.024.

3. Erdos, G., and Dosztányi, Z. (2024). AIUPred: Combining energy estimation with deep learning for the enhanced prediction of protein disorder. Nucleic Acids Res. 52, W176–W181. 10.1093/nar/gkae385.

4. Kliszczak, M., Sedlackova, H., Pitchai, G.P., Streicher, W.W., Krejci, L., and Hickson, I.D. (2015). Interaction of RECQ4 and MCM10 is important for efficient DNA replication origin firing in human cells. Oncotarget 6, 40464–40479. 10.18632/oncotarget.6342.

5. Sievers, F., and Higgins, D.G. (2018). Clustal Omega for making accurate alignments of many protein sequences. Protein Sci. 27, 135–145. 10.1002/pro.3290.

## SUPPLEMENTARY INFORMATION REFERENCES

1. Berger, S., and Chistol, G. (2024). Visualizing the dynamics of DNA replication and repair at the single-molecule level. In Methods in Cell Biology (Elsevier Inc.), pp. 109–165. 10.1016/bs.mcb.2023.07.001.

2. Lebofsky, R., Takahashi, T., and Walter, J.C. (2009). DNA replication in nucleus-free Xenopus egg extracts. Methods Mol. Biol. 521, 229–252. 10.1007/978-1-60327-815-7_13.

3. Sparks, J., and Walter, J.C. (2019). Extracts for analysis of DNA replication in a nucleus-free system. Cold Spring Harb. Protoc. 2019, 194–206. 10.1101/pdb.prot097154.

4. Sparks, J.L., Chistol, G., Gao, A.O., Räschle, M., Larsen, N.B., Mann, M., Duxin, J.P., and Walter, J.C. (2019). The CMG Helicase Bypasses DNA-Protein Cross-Links to Facilitate Their Repair. Cell 176, 167–181.e21. 10.1016/j.cell.2018.10.053.

5. Yardimci, H., Loveland, A.B., van Oijen, A.M., and Walter, J.C. (2012). Single-molecule analysis of DNA replication in Xenopus egg extracts. Methods 57, 179–186. 10.1016/j.ymeth.2012.03.033.

6. Trowitzsch, S., Bieniossek, C., Nie, Y., Garzoni, F., and Berger, I. (2010). New baculovirus expression tools for recombinant protein complex production. J. Struct. Biol. 172, 45–54. 10.1016/j.jsb.2010.02.010.

7. Low, E., Chistol, G., Zaher, M.S., Kochenova, O. V., and Walter, J.C. (2020). The DNA replication fork suppresses CMG unloading from chromatin before termination. Genes Dev. 34, 1534–1545. 10.1101/gad.339739.120.

8. Wu, R.A., Semlow, D.R., Kamimae-Lanning, A.N., Kochenova, O. V., Chistol, G., Hodskinson, M.R., Amunugama, R., Sparks, J.L., Wang, M., Deng, L., et al. (2019). TRAIP is a master regulator of DNA interstrand crosslink repair. Nature 567, 267–272. 10.1038/s41586-019-1002-0.

9. Terui, R., Berger, S.E., Sambel, L.A., Song, D., and Chistol, G. (2024). Single-molecule imaging reveals the mechanism of bidirectional replication initiation in metazoa. Cell 187, 3992–4009.e25. 10.1016/j.cell.2024.05.024.

10. Berger, S., and Chistol, G. (2023). Visualizing the Dynamics of DNA Replication and Repair at the Single-Molecule Molecule Level 1st ed. (Elsevier Inc.) 10.1016/bs.mcb.2023.07.001.

11. Arias, E.E., and Walter, J.C. (2005). Replication-dependent destruction of Cdt1 limits DNA replication to a single round per cell cycle in Xenopus egg extracts. Genes Dev. 19, 114–126. 10.1101/gad.1255805.

12. Sergé, A., Bertaux, N., Rigneault, H., and Marguet, D. (2008). Dynamic multiple-target tracing to probe spatiotemporal cartography of cell membranes. Nat. Methods 5, 687–694. 10.1038/nmeth.1233.

13. Loveland, A.B., Habuchi, S., Walter, J.C., and Van Oijen, A.M. (2012). A general approach to break the concentration barrier in single-molecule imaging. Nat. Methods 9, 987–992. 10.1038/nmeth.2174.

